# Field Plant Biodiversity and Carbon Farming Shape Leaf Microbiomes

**DOI:** 10.64898/2026.09.25.754310

**Authors:** J-B. Floc’h, J. Näsäkkälä, H. Susi, A-L. Laine

## Abstract

Carbon farming practices are increasingly adopted as part of regenerative agriculture to enhance soil carbon sequestration and improve agricultural sustainability. However, their effects on the phyllosphere microbiome, and how these interact with field biodiversity, remain poorly understood. We investigated how field biodiversity and carbon farming practices (cover crops, all- in mixes, adaptive grazing and ley mixture) influence leaf microbial communities across 19 Finnish farms using ITS and 16S amplicon sequencing.

Foliar fungal—but not bacterial— diversity increased with plant species richness. Carbon farming altered both fungal and bacterial community composition, but only fungal community was positively associated with disease load. Network analysis identified three candidate keystone taxa: *Alternaria* sp. (FASV 34) and *Sporobolomyces roseus* (FASV 2070), which were mutually exclusive, and *Methylobacterium* sp. (BASV 75), which co-occurred with FASV 2070.

Together, these findings demonstrate that different carbon farming practices shape distinct leaf microbiomes cover crop systems tending to favor potential fungal pathogens, whereas ley mixture and adaptive grazing strategies promote microbial taxa associated with healthier phyllosphere communities. Our study underscores the importance of understanding the complex microbial interactions within leaf ecosystems to inform sustainable agricultural practices.

## INTRODUCTION

Climate change is reshaping the future of agriculture. Shifts in climate and weather patterns can lead to reduced crop yields due to more frequent and intense extreme weather events, prolonged droughts, unstable winter conditions, increased pest activity, and the spread of invasive species (Cogato et al. 2019; Graczyk and Szwed 2020). Agriculture itself is also a major contributor to climate change, with agri-food systems responsible for an average of 17 billion tons of CO₂ emissions annually (Holka, Kowalska, and Jakubowska 2022). The significant impact of agroecosystems on the biosphere has driven the development of strategies to mitigate greenhouse gas emissions, with a strong emphasis on increased carbon sequestration and storage in agricultural soils and plant biomass (Tiefenbacher et al. 2021; Mattila et al. 2022; Lal 2004). Carbon farming often implemented as part of regenerative farming have been increasingly adopted worldwide (Chenu et al. 2018; Bradford et al. 2019). Carbon farming methods can generally be divided into two categories: those that increase carbon inputs by extending vegetation cover (e.g., longer cropping seasons, higher plant density, adaptive grazing) and increasing plant diversity (e.g., use of cover crops, intercropping, species mixtures), and those that reduce carbon losses by reducing soil disturbance (e.g., no-till cultivation) and improving soil organic matter content (e.g., organic amendments, perennial cropping) (Mattila et al. 2022; Paustian et al. 2019; Teague and Kreuter 2020).

While much is known about how these practices shape soil microbial communities (Bhattacharyya et al. 2022; Clemmensen et al. 2015; Mattila and Vihanto 2024; Domeignoz-Horta et al. 2024), a key challenge is to understand their broader impacts on field-scale ecosystems, including plant diversity and aboveground microbiomes. Carbon farming practices may influence leaf microbiomes through several, potentially interacting mechanisms. First, a more diverse plant community is often associated with increased microbial diversity, as different plant species provide a greater diversity of host traits, organic matter inputs, and habitat niches that can directly influence microbial community assembly (Liebman and Schulte 2015; Matsushita, Yamane, and Asano 2016; Shen et al. 2021; Labouyrie et al. 2023). More diverse plant assemblages are also associated with greater network connectivity in microbiome network structures (Teague and Kreuter 2020; Domeignoz-Horta et al. 2024). Second, carbon farming practices may influence the microbiome through changes in soil conditions (Cesarano et al. 2017). Soil amendments (nutrient-rich, e.g., manure; nutrient-poor, e.g., biochar, paper mill sludge) and reduced soil disruption (e.g., no-till) can alter soil physicochemical properties, structure, and microbial communities (Cesarano et al. 2017). These belowground changes may, in turn, shape the foliar microbiome, because part of the leaf microbiome originates from the soil, with this influence stronger for fungi than for bacteria (Hamonts et al. 2018). Third, cattle grazing can alter host– pathogen–herbivore interactions, thereby affecting plant disease dynamics and leaf microbial communities (Li et al. 2024; Borer et al. 2009). Finally, the diversity leaf microbial communities may also be influenced by field remnants, such as patches of semi-natural vegetation within agricultural landscapes, that may serve as reservoirs of microbial taxa that buffer crops against disturbances (Smith et al. 2018; Alarcón-Segura et al. 2022; Holka, Kowalska, and Jakubowska 2022). However, the extent to which these mechanisms shape leaf microbial communities remains poorly understood.

The leaf microbiome represents a critical but understudied component of agroecosystems. Leaf-associated bacteria and fungi play an essential role in plant health (Beattie and Lindow 1999; Vacher et al. 2016; Ritpitakphong et al. 2016). Both fungal and bacterial endophytes can directly influence crop yield and promote plant growth (Yashaswini, Nysanth, and Anith 2021; Akköprü et al. 2021; Poveda et al. 2019). Leaf endophytes also play a crucial role in pathogen suppression. For instance, fungal endophytes secrete antibiotics that suppress pathogenic fungi responsible for leaf diseases (Grabka et al. 2022; González-Teuber et al. 2021). Experimental studies have also shown that cover crops can suppress the foliar pathogen *Pseudomonas syringae* (Maglione et al. 2024). Because leaves host both resident and transient microbial taxa and integrate microbial inputs from both the surrounding environment and belowground compartments, the phyllosphere provides an ideal system for evaluating how carbon farming practices and plant diversity jointly shape microbial communities across the entire plant holobiont, linking aboveground microbial communities with belowground root and soil microbiomes. This connection is especially evident in leaf endophytic communities, as certain endophytes can migrate from the roots to the leaves, establishing a dynamic interaction between belowground and aboveground microbial communities (Chi et al. 2005; Wu et al. 2025; Pangesti et al. 2020). By investigating leaf microbiomes across farms differing in carbon farming practices and biodiversity, we aim to clarify how these practices restructure aboveground microbial networks and their implications for crop health and resilience.

Unraveling the complexity of leaf-associated microbiomes requires high-resolution tools capable of capturing microbial diversity and community composition. Next-generation sequencing (NGS) has revolutionized microbial ecology by enabling culture-independent characterization of entire microbial communities with unprecedented depth and accuracy (Garg et al. 2024; Ronholm 2018). Amplicon sequencing, targeting marker genes such as the 16S rRNA gene for bacteria and the ITS (Internal Transcribed Spacer) region for fungi, has become the standard approach for characterizing microbial community composition across diverse environments (Tedersoo and Lindahl 2016; Callahan et al. 2019; Baldrian 2019). These sequencing techniques enable the assessment of microbial diversity and community shifts in response to environmental factors, including agricultural management practices (Mattila and Vihanto 2024; Floc’h et al. 2020).

In this study, we investigated how carbon farming practices and field remnant biodiversity influence the microbial diversity and composition of leaf microbial communities in crops and remnant vegetation across Finnish farms. We also qualified disease symptoms to examine their associations with leaf microbial communities. We sampled crops and remnant vegetation across 19 Finnish farms representing four carbon farming practices. Specifically, we tested:

1) Does alpha diversity of microbial communities of carbon farmed and control farms vs remnants differ and do we see correlation between microbial diversity and plant diversity in the different treatments?
2) Does microbiome community structure, occurrence network and keystone species differ among carbon farming treatments, their control and remnants?
3) Do carbon farming practices alter disease load in plants and can we find associations between specific herbivory and disease symptoms and microbes?

We found that the foliar fungal—but not bacterial—diversity increased with plant species richness. Carbon farming practices altered both fungal and bacterial community composition, but only fungal communities were positively associated with disease load. Network analysis highlighted three potential keystone taxa: *Alternaria* sp. (FASV 34) and *Sporobolomyces roseus* (FASV 2070), which were mutually exclusive, and *Methylobacterium* sp. (BASV 75), which co-occurred with FASV 2070. These results suggest that carbon farming practices shape distinct leaf microbiomes: cover crops tended to favor fungal pathogens, whereas ley mixture and adaptive grazing supported beneficial taxa. Together, these findings demonstrate that carbon farming practices shape distinct phyllosphere microbiomes and provide a foundation for developing microbiome-based strategies to enhance crop resilience.

## MATERIALS AND METHODS

### 1. Sampling sites and vegetation survey

The Carbon Action experiment, coordinated by the Baltic Sea Action Group (Espoo, Finland), was launched in 2018 as a five-year study involving 105 farms across Finland to investigate how carbon farming practices influence soil carbon sequestration (Mattila et al. 2022). The farms comprised both pastures and agricultural croplands, with field locations initially identified using satellite imagery (Nevalainen et al. 2022). For the present study, we selected a subset of 19 farms located in Southern and Central Finland. The selected farms represented four carbon farming practices: undersown cover crops with the main crop (caraway, pea, rye, or oats; hereafter referred to as cover crops, 6 farms), ley mixtures (5 farms), adaptive grazing designed to enhance carbon sequestration (hereafter adaptive grazing, 4 farms), and an “all-in” practice (4 farms). Since 2019, each farm had implemented one of the carbon farming practices in a 1.5-hectare field plot (Figure 1). Adjacent control plots were managed using conventional practices for the same crop or pasture. The all-in treatment combined undersown cover crops with locally adapted soil management measures, including no tillage and subsoiling. Its corresponding control consisted of the same crop but without cover crops or soil amendments. Adaptive grazing consisted of high stocking density, frequent rotation, and extended recovery periods for vegetation (Cappelli et al. 2024), while the control treatment used continuous grazing. Both adaptive grazing treatments and controls were based on mixtures of gramineous and leguminous forage species. For the cover crop treatment, the control involved growing the same main crop (caraway, pea, rye, or oats) without undersowing. The control ley mixture contained no more than three sown species. Cover crop, all-in, and ley mixture treatments included a greater number of sown species than their respective controls. Previous studies have shown that adaptive grazing can influence plant species richness, although reported effects vary among systems (Li et al. 2024; Morris 2021).

**Figure 1.**
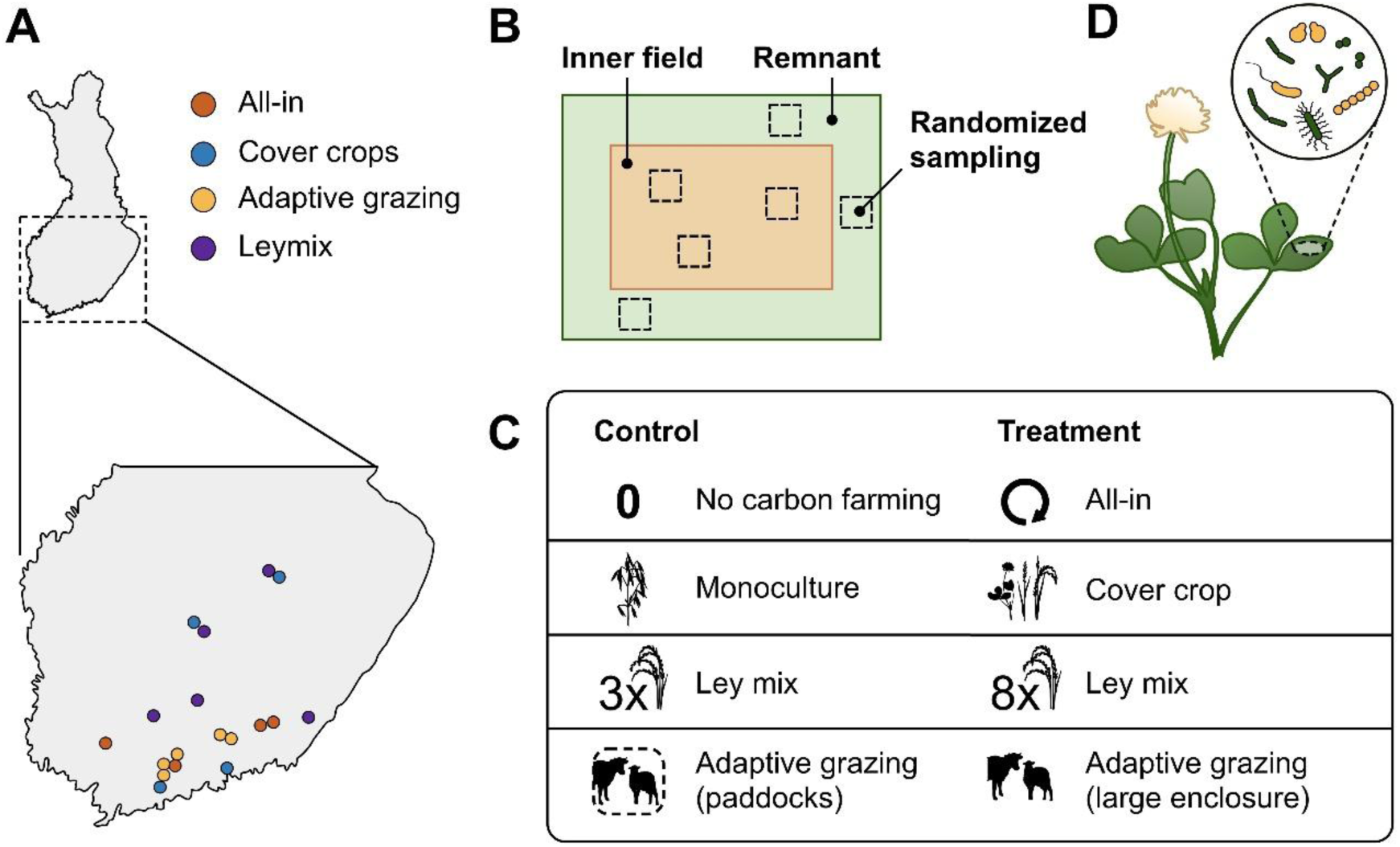
Sampling design of our experiment. The leaf samples were taken from randomly chosen plots (1 m^2^) in the remnants (light green) and in the field (light orange).

We conducted vegetation surveys and sampling within 12 days’ time on 26 June – 7 July 2023 and at each farm we sampled both the carbon farming treatment and its corresponding control plot. We placed a 1 m^2^ quadrats in carbon-farmed fields, control fields, and field remnants to assess vegetation richness and diversity. Quadrat locations were randomized by tossing a tennis ball and placing the quadrat where it landed. At each farm, twelve 1 m^2^ quadrats were surveyed including six from carbon-farmed fields and six from control fields, together with remnant patches – resulting in a total of 228 vegetation quadrats. From both the interiors and remnants of these fields, we identified plant species, estimated their cover, and recorded symptoms of plant diseases and herbivory. From each field plot, six crop plant samples and six cover crop plants were sampled for symptom assessment. In addition, six wild plant individuals were sampled from the remnant patches. For all groups, the most common plant species were selected for sampling. Individual plants were chosen randomly, irrespective of symptom presence. Because disease symptoms and herbivore damage frequently co-occur (Nakazawa, Yamanaka, and Urano 2012), both were assessed as the percentage of leaf area exhibiting characteristic herbivory symptoms (large holes, small holes, window feeding, thrips, aphids, leaf miners, scraping, moth damage, spittle, mites, galls; caterpillar see Supplementary Table S1 for explanation) and disease symptoms (rust, mildew, leaf spot, virus symptoms; see Supplementary Table S1 for explanation). Assessments were conducted on the five oldest leaves of each plant. To characterize the microbiome of the scored plants, we collected leaf material for subsequent DNA extraction and analysis. Plants were sampled irrespective of visible disease symptoms. We excised 1.5 -cm leaf sections using forceps disinfected between samples with DNA Away® and milli-Q® water and stored in 2 mL Eppendorf tubes. In total, 293 unique leaf samples were included in the microbial community analyses, of which 207 were analyzed for fungal (ITS) communities and 229 for bacterial (16S) communities, with 143 samples shared between the two datasets.

### 2. DNA extraction

DNA was extracted following the protocol of Lodhi et al. (1994. The extraction buffer included 2.0 % (w/v) cetrimonium bromide (CTAB), 100 mM tris-HCl, and two drops of 0.2% β-mercaptoethanol per preparation. In addition, 50 mg of polyvinylpolypyrrolidone (PVP) was added to 15 mL of buffer, and NaCl. The full extraction process can be found in the supplementary materials. After extraction, samples were stored in −20 °C until further analysis.

### 3. Library preparation, PCR amplification and Sequencing

Extracted DNA samples were sent to BioName (Turku, Finland) for PCR amplification and sequencing. Fungal ITS2 and bacterial 16S rRNA regions were amplified using the primer pairs: fITS7 (GTGARTCATCGAATCTTTG) and ITS4 (TCCTCCGCTTATTGATATGC) for the fungal ITS2 region (Ihrmark et al. 2012) and Bakt_341F (CCTACGGGNGGCWGCAG) and Bakt_805R (GACTACHVGGGTATCTAATCC) (Herlemann et al. 2011). All primers were modified with linker tags and heterogeneity spacers to enable attachment of NGS adapters and improve amplicon library diversity. Each sample was amplified in two technical replicates, and each replicate included two heterogeneity spacer variants of each primer. For fungal ITS2 libraries, only samples yielding sufficient PCR amplification were selected for sequencing, whereas no such pre-selection was required for bacterial 16S libraries. Library preparation followed BioName’s internal dual-indexing protocol, which included unique combination of i5 and i7 indices across replicates. PCR products were pooled and purified using the Genejet gel extraction kit (Thermo Scientific). Sequencing was performed on an Illumina NovaSeq 6000 platform with paired-end 2 x 250 bp reads at the Finnish Functional Genomic Centre.

#### Bioinformatic processing

The following bioinformatic analysis was performed after BioName’s pipeline. Paired-end reads were merged using VSEARCH v2.22.1 (Rognes et al. 2016) and primer trimming was performed with CUTADAPT v3.5 (Martin 2011), allowing up to 20% mismatches and length filters specific to each marker region (ITS2: 100-1000 bp, 16S: 377-477 bp). Reads were quality-filtered with a maximum of one expected error and dereplicated by removing singletons. Denoising and generation of zero-radius operational taxonomic units (ZOTUs) were conducted using the UNOISE3 algorithm in USEARCH v11 (Nilsen et al. 2024; Edgar 2010). After filtering for chimeras, index cross-talk (tag-jumps), and low-abundance features (< 2 reads), we proceeded with taxonomic classification independently from Bioname. Taxonomy was assigned using the RDP classifier v2.14 (Wang and Cole 2024) with the UNITE reference database (Kõljalg et al. 2005) for fungal ITS2 and the SILVA v138 database (Quast et al. 2013) for bacterial 16S sequences. Final datasets included both absolute and relative abundance for downstream ecological analyses.

### 4. Statistical and network analysis of microbial community data

All statistical analyses were conducted in R version 4.4.0 (Core Team 2013) using Rstudio version 2025.03.0+400 (Posit Software, PBC 2025). First, to understand how fungal and bacterial alpha diversity varies among carbon practices and their controls, fungal and bacterial alpha diversity was quantified for each plant using Shannon’s diversity index (Shannon 1948). plant species diversity (Shannon’s) was calculated for each field plot to examine its relationship with microbial diversity. Shannon’s index was selected because it incorporates both species richness and evenness and is widely used in microbial and ecological studies (Feranchuk et al. 2018). Shannon’s index was calculated using the *vegan* package v2.6-10 (Oksanen et al. 2024) as follows:

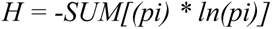

where *pi* is proportional abundance of species *i.* Microbial Shannon diversity was calculated separately for each sampled plant, whereas plant Shannon diversity was calculated at the field-plot level. For analysis relating plant and microbial diversity, each microbial sample was assigned the plant Shannon diversity value of the field plot from which it originated. within each plant (microbial communities) or field plot (plant communities). Linear models were fitted in R v4.4.0 to test the relationship between plant and microbial alpha diversity and whether this relationship differed among carbon farming treatments. [plant/microbial] Shannon diversity was used as the response variable, with [microbial/plant] Shannon diversity, carbon farming treatment, and their interaction as explanatory variables. Post-hoc comparisons were performed using *emmeans* v1.11.1 (Lenth 2024) with compact-letter displays generated using *multcompView* v0.1-10 (Spencer Graves 2024).

Secondly, we investigated whether microbial community composition, co-occurrence network structure, and keystone taxa differed among carbon farming treatments, their corresponding controls, and field remnants. Differences in microbial community composition were assessed using permutational multivariate analysis of variance (PERMANOVA) implemented in the *vegan* package. Fungal and bacterial community dissimilarity matrices were calculated separately from the ITS and 16S abundance tables, respectively, using Bray–Curtis dissimilarities. PERMANOVA models included carbon farming treatment, sampling location (field interior or remnant vegetation), host plant species, and their interactions as explanatory variables. To account for the sampling design and non-independence of samples from the same farm, permutations were constrained by farm identity. Pairwise PERMANOVA comparisons were subsequently performed using the *pairwiseAdonis* package v0.4 ((Martinez Arbizu 2020).

To test whether carbon farming practices favor certain microbial taxa, we performed indicator species analysis using the multipatt function from the indicspecies package v1.8.0 (Cáceres and Legendre 2009). In this analysis, carbon farming treatments and their corresponding controls were used as grouping variables. Separate analyses were run for fungi and bacteria. To further explore how carbon farming practices and sampling location (field interior or remnant vegetation) shaped the plant leaf microbiome, we constructed co-occurrence networks at two levels: between fungal and bacterial communities (interkingdom) and within each kingdom. These networks were used to identify potential keystone taxa within each experimental condition and spatial location. Microbial co-occurrence networks were inferred using SPIEC-EASI v1.1.3 (Kurtz et al. 2015) with the Meinhausen-Bühlmann (MB) algorithm, and network visualizations were created in Cytoscape v3.10.3 (P. Shannon et al. 2003). Statistical plots were produced using ggplot2 v3.5.1 (Wickham 2016).

To test whether certain microbial taxa are associated with herbivory and disease symptoms in the plants, we performed indicator species analysis using the multipatt function from the indicspecies package v1.8.0 (Cáceres and Legendre 2009). In this analysis, we used the different symptom catergories (Table S1) as grouping variables. Separate analyses were performed for fungal and bacterial communities. Finally, to disentangle direct and indirect effects of environmental factors (carbon farming treatment, farm identity, and sampling location (field interior or remnant vegetation)) on microbial community composition and disease load, we applied partial least squares path modeling (PLS-PM). Because of the limited sample size, we did not perform bootstrapping to estimate the statistical significance of path coefficients. Bootstrap resampling in PLS-PM can produce unstable estimates and inflated standard errors when sample sizes are limited. Instead, model interpretation focused on direction and magnitude of the path coefficients, following recommendations PLS-PM analyses for based on limited sample sizes (Hair et al. 2022).

## RESULTS

### 1. Alpha diversity of the bacterial and fungal leaf communities

We first assessed the effects of carbon farming practices on the alpha diversity of leaf microbial communities. Carbon farming practices significantly affected both bacterial (*F* = 2.877, *p* = 0.017) and fungal alpha diversity (*F* = 3.526, *p* = 0.003; Table 1). In contrast, the effect of host plant identity was only marginally significant for bacterial alpha diversity (*p* = 0.078) and was not significant for fungal alpha diversity (*p* = 0.12).

**Table 1.**
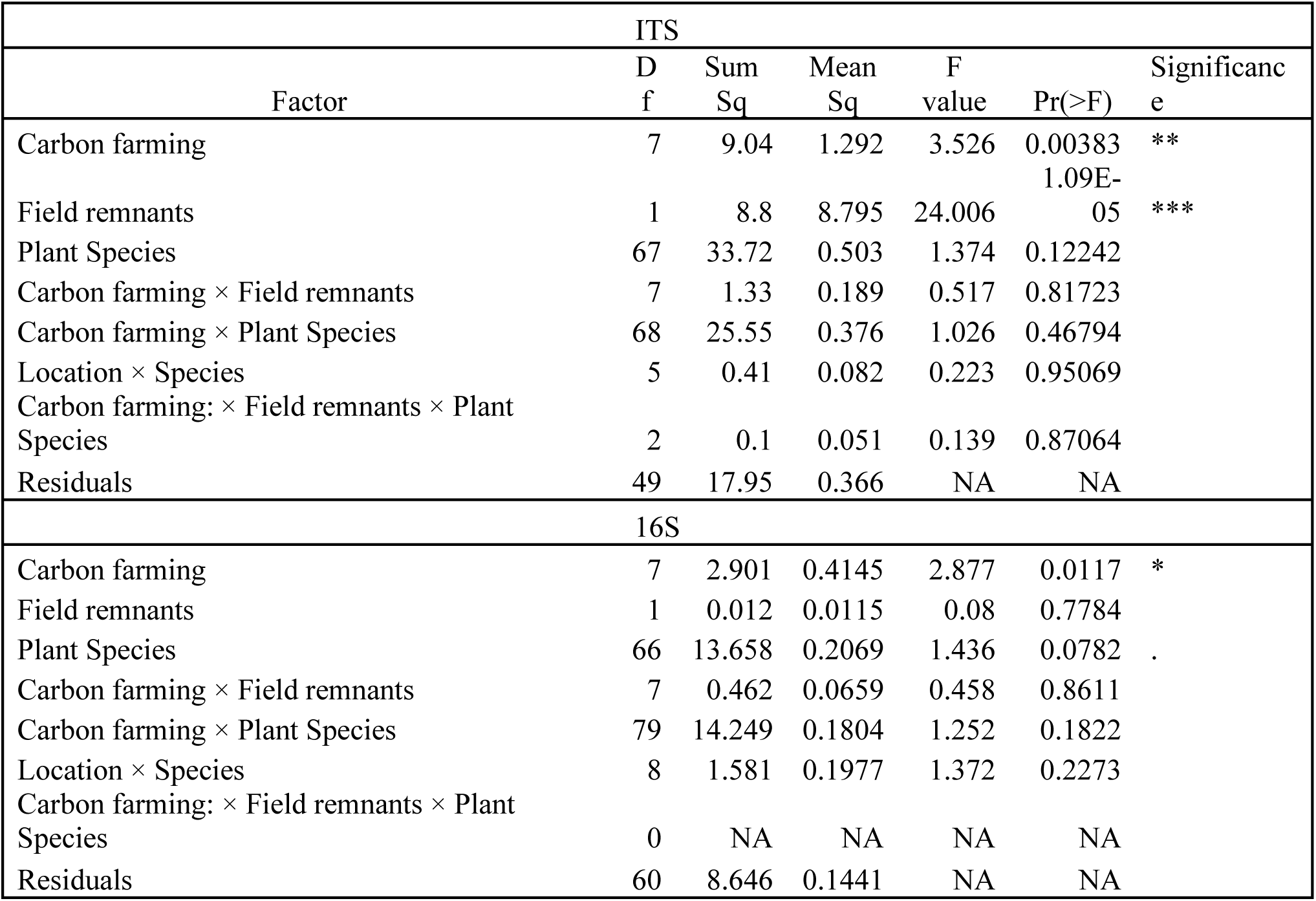
Effect of the carbon farming treatments on the alpha diversity of the leaf microbiome (Shannon index) according to ANOVA, n_ITS_ = 207 and n_16S_ = 229, α = 0.05, Significance codes: p = 0 ‘***’ p = 0.001 ‘**’ p = 0.01 ‘*’ p = 0.05 ‘.’ p = 0.1 ‘ ’ p = 1.

Sampling location (field interior or remnant vegetation), used as a proxy for plant community context, had no significant effect on bacterial alpha diversity (p = 0.78), but had a strong positive effect on fungal alpha diversity (p < 0.001) explaining a substantial proportion of the variation in the ITS dataset.

Linear regression analyses revealed positive relationships between plant diversity and fungal alpha diversity (Figure 2). In particular, the ley mixture and adaptive grazing treatments exhibited significant positive relationships between plant and fungal diversity. Although no significant relationships were detected for the remaining treatments, regression slopes were predominantly positive, suggesting a general tendency for greater plant richness to be associated with higher fungal diversity in the phyllosphere.

**Figure 2.**
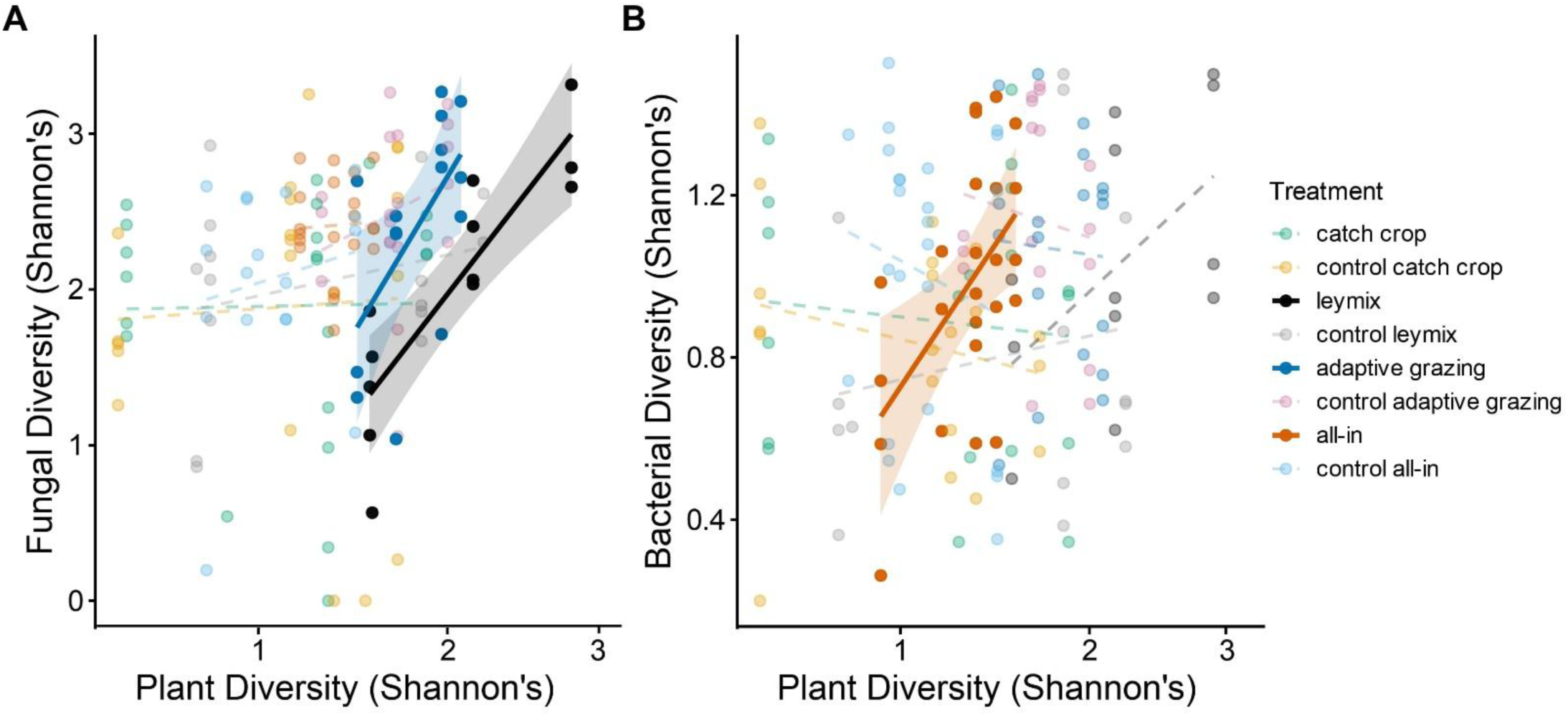
Correlation between plant and fungal diversity, and between plant and bacterial diversity, in the plant leaf microbiome for each carbon farming practice, according to linear regression models (lmer), nITS = 207, n16S = 229, α = 0.05.

In contrast, the relationships between plant and bacterial diversity were weaker and more variable than those observed for fungi (Figure 2). Among all carbon farming practices, only the all-in treatment showed a strong and significant positive correlation between plant and bacterial alpha diversity. Most remaining treatments showed negative but non-significant slopes, indicating no consistent relationship between plant and bacterial diversity under those management conditions.

### 2. Microbial community dynamics

#### a. Carbon farming significantly affects the structure of plant leaf microbial communities

Carbon farming treatments also influenced leaf microbiome structure, as reflected in both the taxonomic profiles (Figure 3) and the PERMANOVA analyses (Table 2). Overall, both bacterial and fungal communities were dominated by a relatively small number of taxonomic families across all carbon farming practices, although their relative abundances varied among treatments and their corresponding controls. In bacterial communities, Sphingomonadaceae represented the most abundant family in nearly all treatments, whereas fungal communities were primarily composed of members of the Tremellales, Pleosporaceae, Phaeosphaeriaceae, and several unclassified Ascomycota. Despite this broadly conserved taxonomic composition, the relative abundance of dominant taxa differed among carbon farming practices, indicating that management altered community structure without changing the dominant microbial groups. For the fungal communities (ITS), host plant identity explained the largest proportion of variation (R^2^ = 0.380, p < 0.001). Carbon farming treatment also had a significant effect (R^2^ = 0.076, p < 0.001), as did sampling location (field interior or remnant vegetation; R^2^ = 0.037, p < 0.001; Table 2). Similarly. for bacterial communities (16S), host plant identity explained the largest proportion of variation (R^2^ = 0.355, p < 0.001). Carbon farming treatments also had significant effect (R^2^ = 0.065, p < 0.001) whereas sampling location explained smaller but proportion of variation (R^2^ = 0.005, p = 0.003). Contrast to the fungal communities, all-interaction terms were significant for the bacterial communities, although they explained only small to moderate to small proportions of the observed of variation (Table 2). This suggests that bacterial communities responded more strongly to interactions among host and environmental factors whereas fungal communities were influenced primarily by the main effects.

**Table 2.**
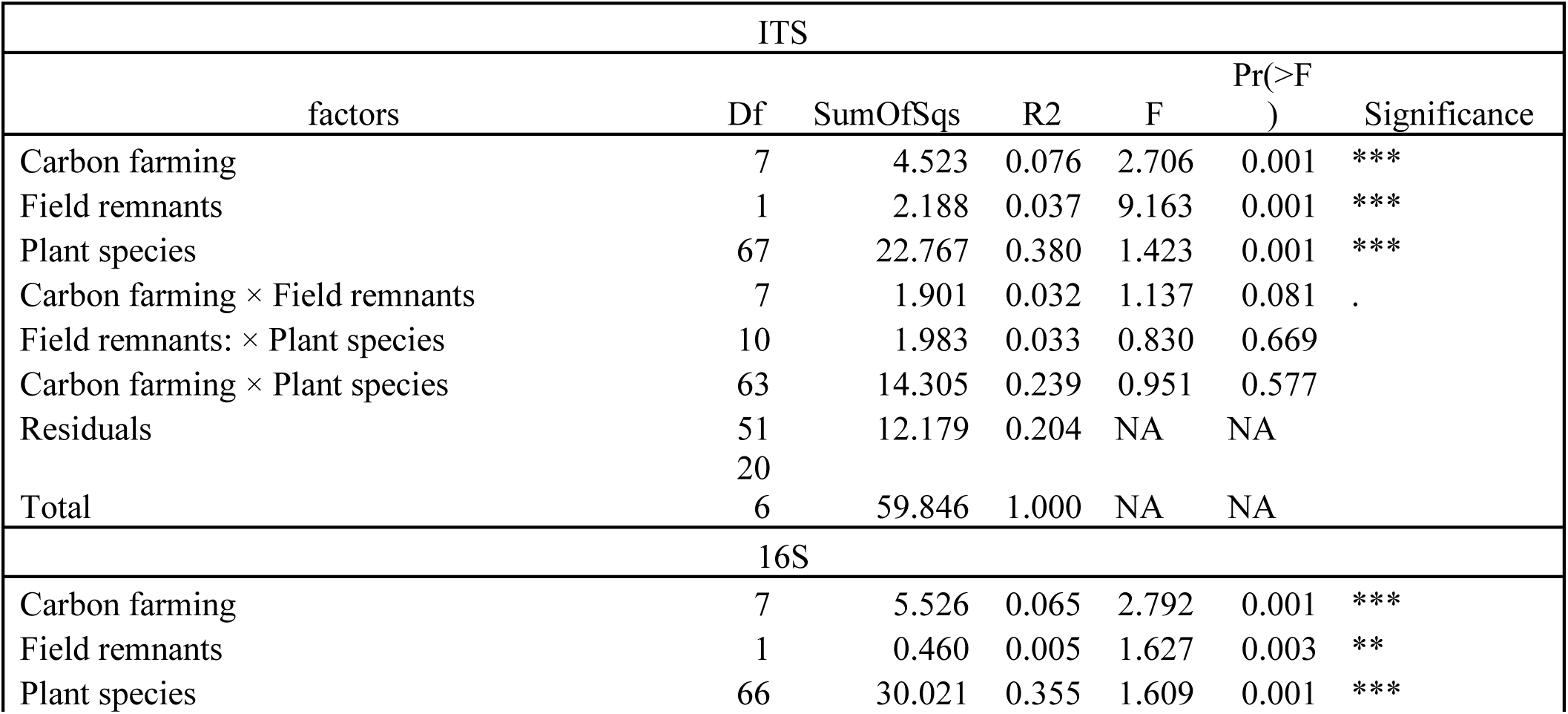
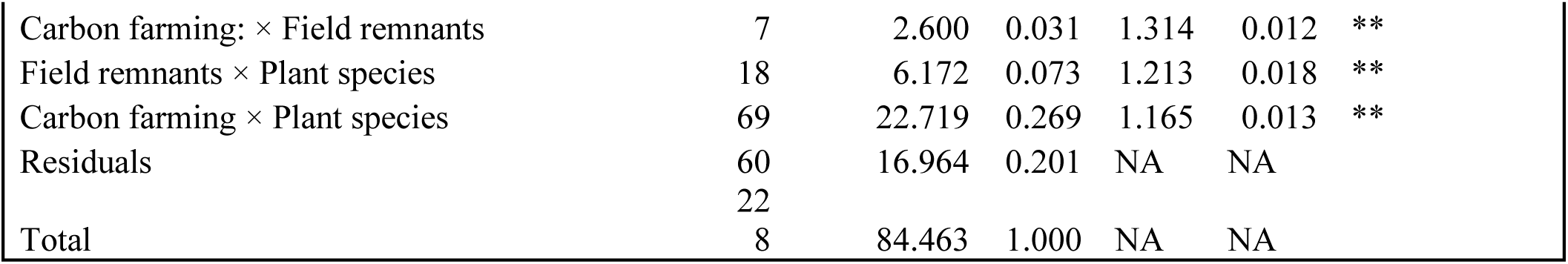
Effect of the carbon farming practices on the structure of the fungal (ITS) and bacterial (16S) communities of the plant leaf microbiome according to PERMANOVA, n_ITS_ = 207 and n_16S_ = 229, α = 0.05, Signif. codes: p = 0 ‘***’ p = 0.001 ‘**’ p = 0.01 ‘*’ p = 0.05 ‘.’ p = 0.1 ‘ ’ p = 1.

**Figure 3.**
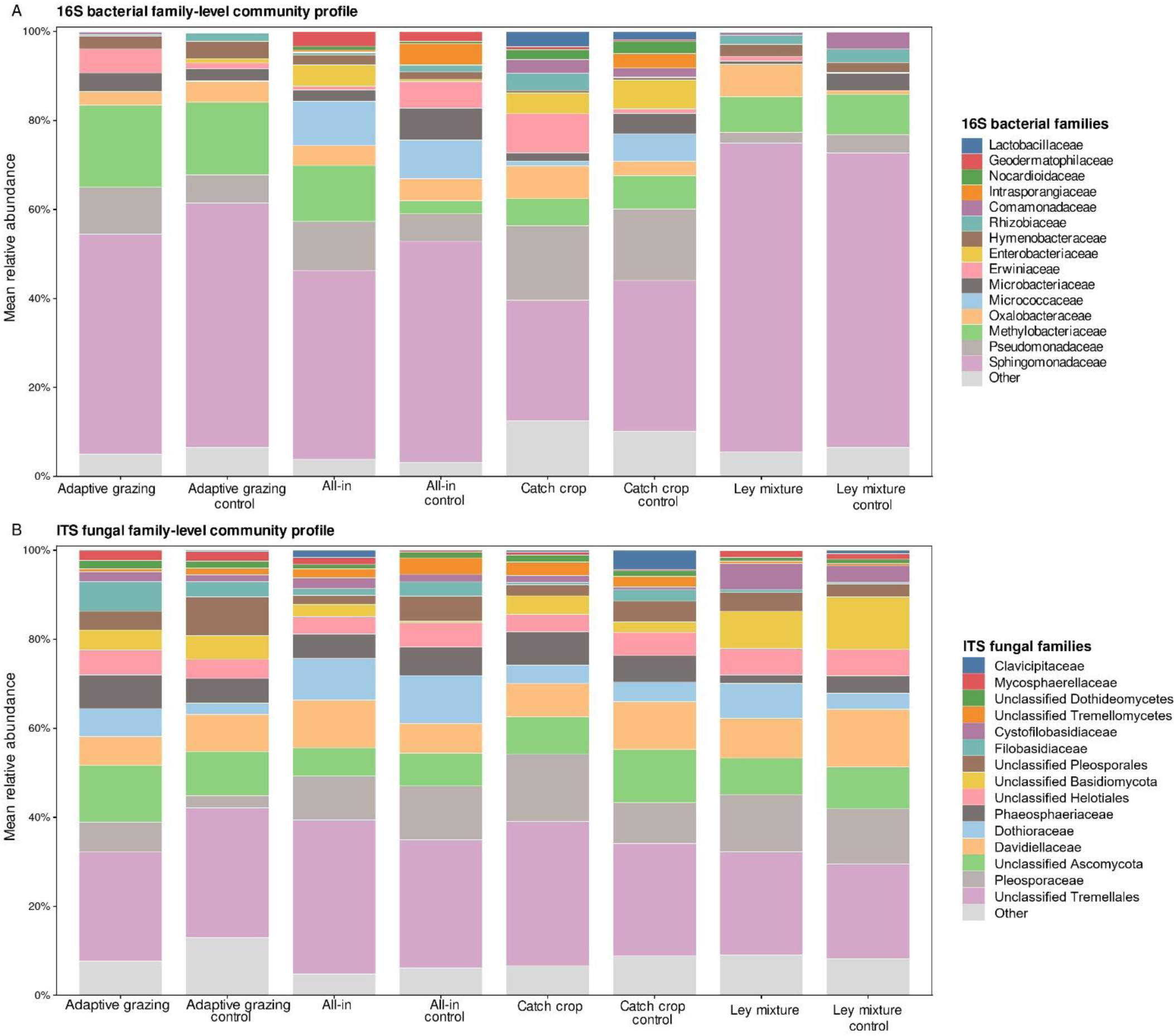
Taxonomic profile of the bacterial (16S) and fungal (ITS) communities in the plant phyllosphere, with and without carbon farming. All the taxa with an abundance of less than 1% of the total community were discarded from the plots.

To identify which specific carbon farming practice contributed to change in microbial community composition, we performed MRPP pairwise comparisons. For fungal communities (Table 3), significant pairwise differences were detected between “all-in” and “ley mixture” (F = 3.30, R^2^ = 0.068, p.adj = 0.028), “cover crop” and “ley mixture” (F = 2.75, R^2^ = 0.054, p.adj = 0.028), “cover crop” and “control ley mixture” (F = 2.74, R^2^ = 0.050, p.adj = 0.028) and finally between “control all-in” and “ley mixture” (F = 3.06, R^2^ = 0.068, p.adj = 0.028). Together with comparisons involving adaptive grazing (Table 3) these results suggest that perennial management practices produced the strongest shifts in fungal community composition. Notably, fungal communities did not differ significantly between most treatments and their respective controls (e.g., “cover crop” vs “control cover crop”, p.adj = 1.000), suggesting that treatment identity, rather than carbon addition alone drives differences in fungal community composition.

**Table 3.**
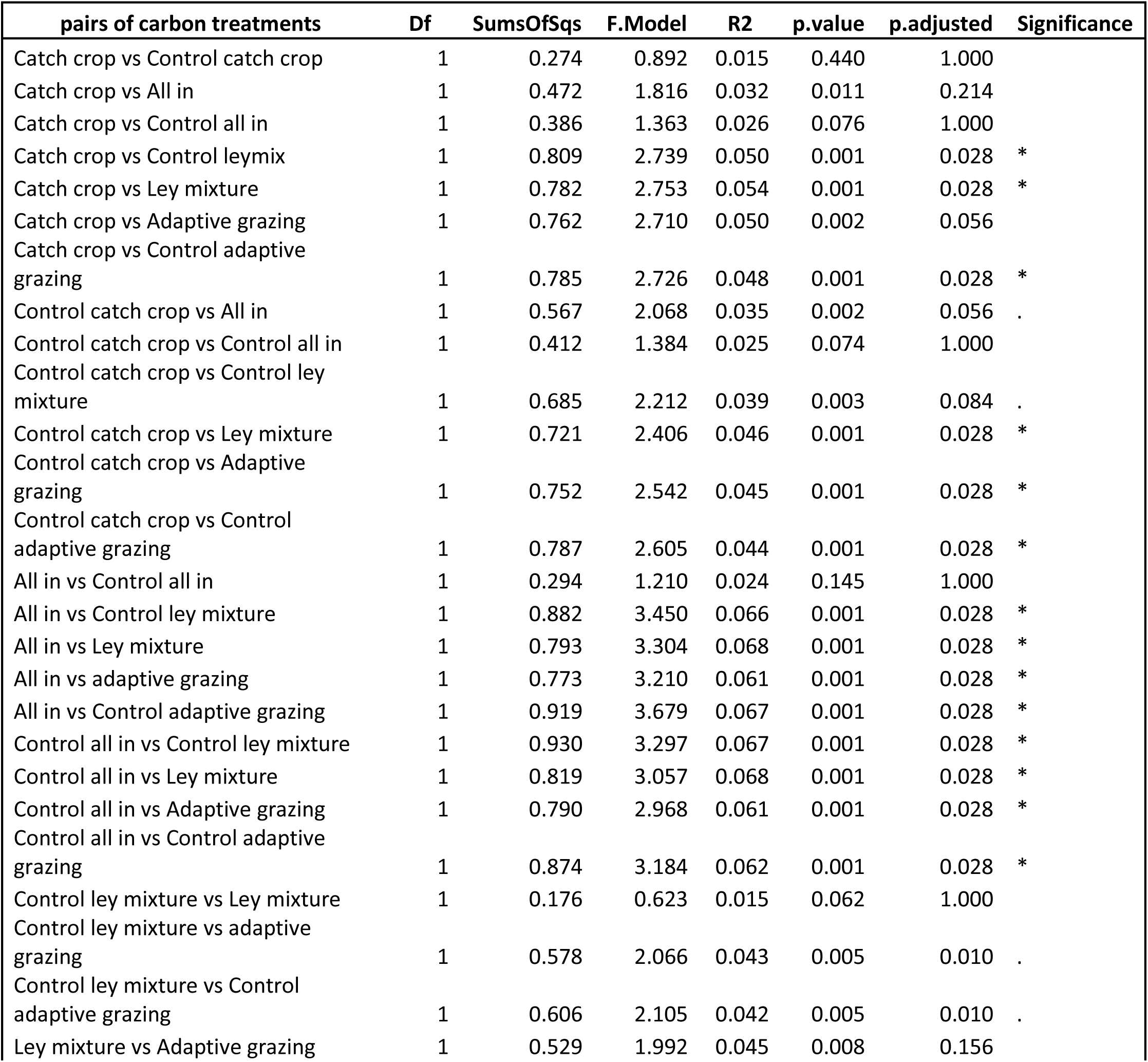
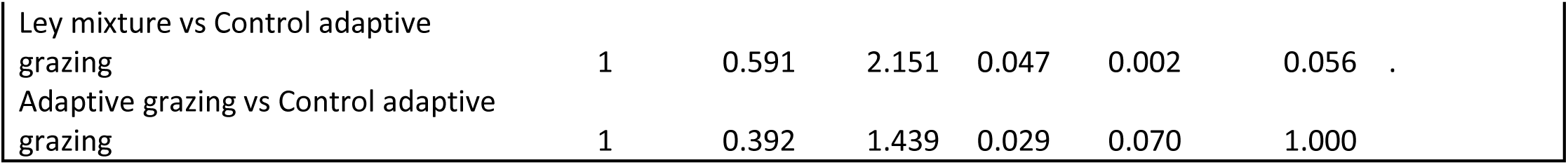
Pairwise comparison of the structure of the fungal communities of the plant leaf microbiome, depending on the carbon farming practices, according to Multi Response Permutation Procedure. α = 0.05, n = 207, Signif. codes: p = 0 ‘***’ p = 0.001 ‘**’ p = 0.01 ‘*’ p = 0.05 ‘.’ p = 0.1 ‘ ’ p = 1.

In the bacterial dataset (Table 4), fewer pariwise comparisons remained significant after correction for multiple testing. Significant differences were detected between “cover crop” and “leymix” (F = 4.94, R^2^ = 0.078, p.adj = 0.028), “control leymix” (F = 4.44, R^2^ = 0.075, p.adj = 0.028) and “control adaptive grazing” (F = 3.71, R^2^ = 0.058, p.adj = 0.028). Although some pairwise comparisons explained relatively large proportions of variation, bacterial communities exhibited fewer significant compositional differences among treatments than fungal communities. As with fungi, comparisons between carbon farming treatments and their corresponding controls were generally not significant.

**Table 4.**
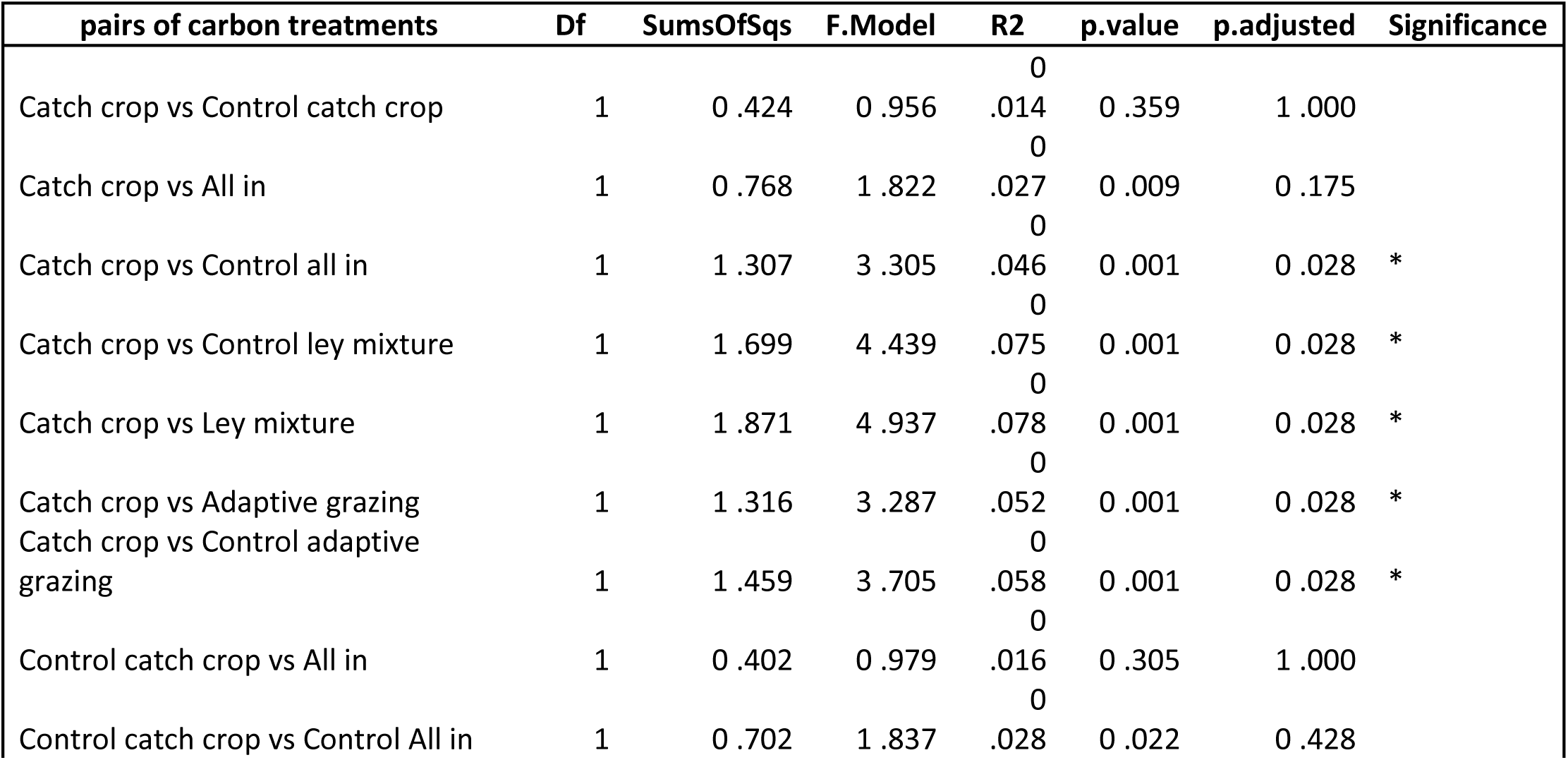
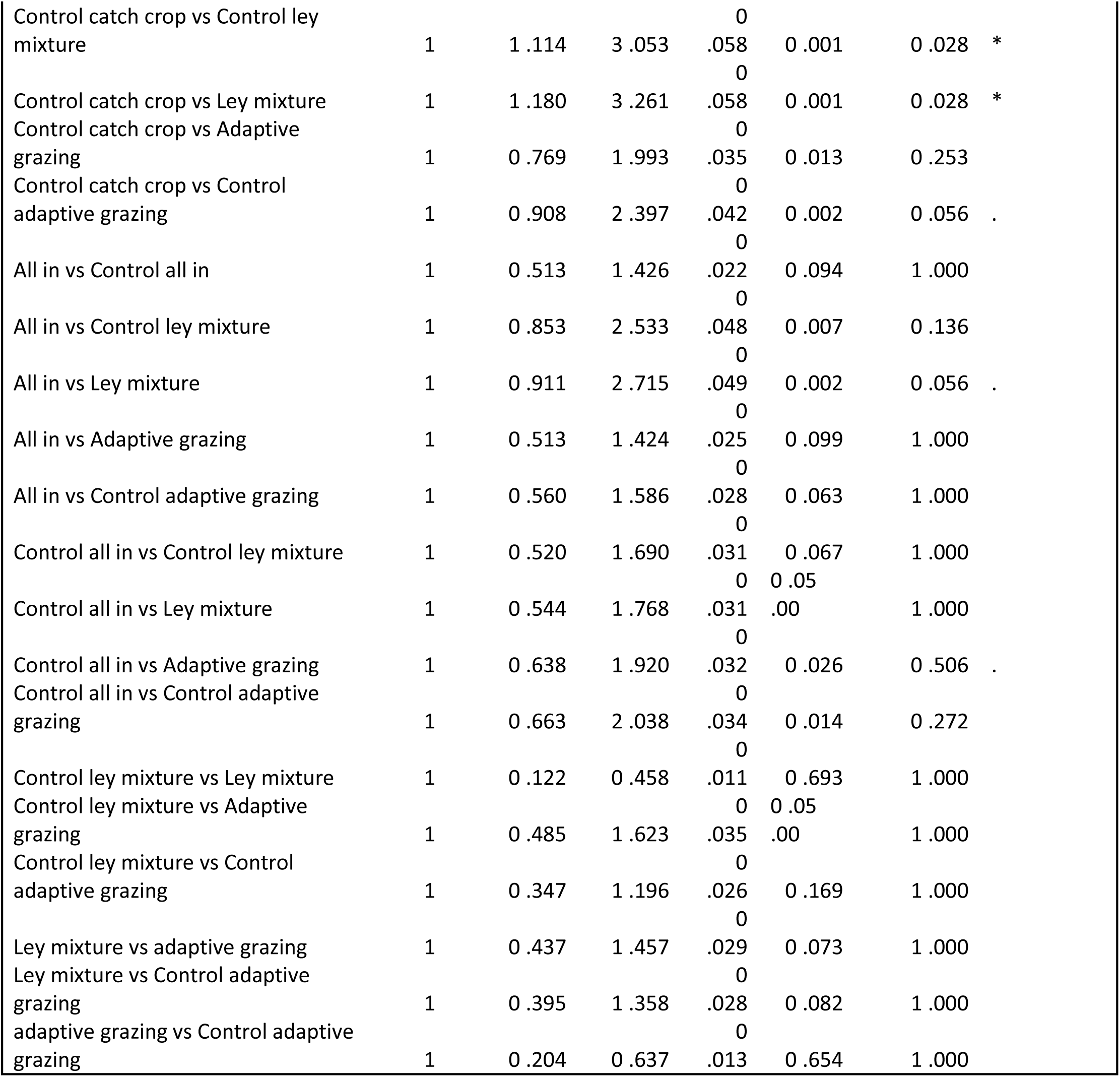
Pairwise comparison of the structure of the bacterial communities of the plant leaf microbiome, depending on the carbon farming practices, according to MRPP. α = 0.05, n = 229, Signif. codes p = 0 ‘***’ p = 0.001 ‘**’ p = 0.01 ‘*’ p = 0.05 ‘.’ p = 0.1 ‘ ’ p = 1.

Together these results demonstrate that carbon farming practices influence the composition of both fungal and bacterial phyllosphere communities. Fungal communities showed more consistent differences among treatments whereas and bacterial communities exhibited fewer significant compositional changes.

##### Carbon farming favorize specific fungi and bacteria

Indicator species analysis revealed control treatments generally contained more fungal and bacterial indicator taxa than the corresponding carbon farming treatments (Supplementary Tables S2–S3).

For fungi (Supplementary Table S2), adaptive grazing contained the highest number of fungal indicator taxa among the carbon farming treatments (18 indicators), dominated by *Ascomycota* and *Phaeosphaeriaceae* members, including *Kabatiella* and *Bulmeria*. Its control contained 13 indicators such as *Sarocladium*, *Myrothecium* and *Phoma*. Control ley mixture contained 10 indicator taxa, including members of *Phaeosphaeria*, *Itersonilia* and *Leptosphaeria*, whereas the ley mixture itself had seven, including unidentified *Exobasidiomycetes* and *Dioszegia*. The all-in treatment and its corresponding control contained three and six fungal indicators, respectively, with the latter dominated by *Cryptococcus* ZOTUs. Control cover crop showed a single *Ascomycota* indicator whereas cover crop treatment had no indicator taxa.

For bacterial communities (Supplementary Table S3), control treatments generally contained more indicator taxa than their corresponding carbon farming treatments. Control ley mixture contained the greatest number of bacterial indicator taxa (40 indicators), including multiple *Hymenobacter, Pedobacter* and *Herbiconiux* taxa. Ley mixture contained seven indicators, dominated by *Sphingomonas*, *Bosea* and *Poladomonas*. Adaptive grazing had five bacterial indicators, including *Sphingomonas*, *Azomonas* and *Teluria*, while its control had 18, such as *Rhodococcus* and Hymenobacter. All-in had a single bacterial indicator (Pseudomonas), while its control had three, including *Oryzihumus*. Cover crop and its control had no indicators.

#### b. Microbial co-occurrence networks and potential keystone taxa

To further explore how experimental conditions shaped the plant leaf microbiome, we constructed co-occurrence networks at two levels: between fungal and bacterial communities (interkingdom) and within each kingdom. These networks were used to identify potential keystone taxa under each experimental condition and spatial location.

Interkingdom networks consistently separated into fungal and bacterial subnetworks under all experimental (Figure 4 and Supplementary Figure S1). Network complexity differed among treatments with control plots forming denser networks than carbon farming plots (189 nodes, 415 edges 126 vs. nodes and 170 edges), suggesting more numerous microbial associations under conventional management. Networks from the field interiors and remnants were similar in size but differed in cross-kingdom connectivity with remnant vegetation exhibiting slightly more bacterial-fungal associations.

Several keystone taxa were repeatedly identified across experimental conditions. (Table 5-7). *Methylobacterium* (Bzotu75) appeared as a keystone in carbon farming, control and field bacterial carbon management networks, suggesting that it represents a potential core bacterial connector across conditions. Likewise, *Sporobolomyces roseus* (Fzotu2070), was consistently identified in interkingdom and fungal networks as a keystone whereas *Alternaria* (Fzotu34) functioned as a keystone taxa under carbon farming and in remnant fungal communities.

**Table 5.**
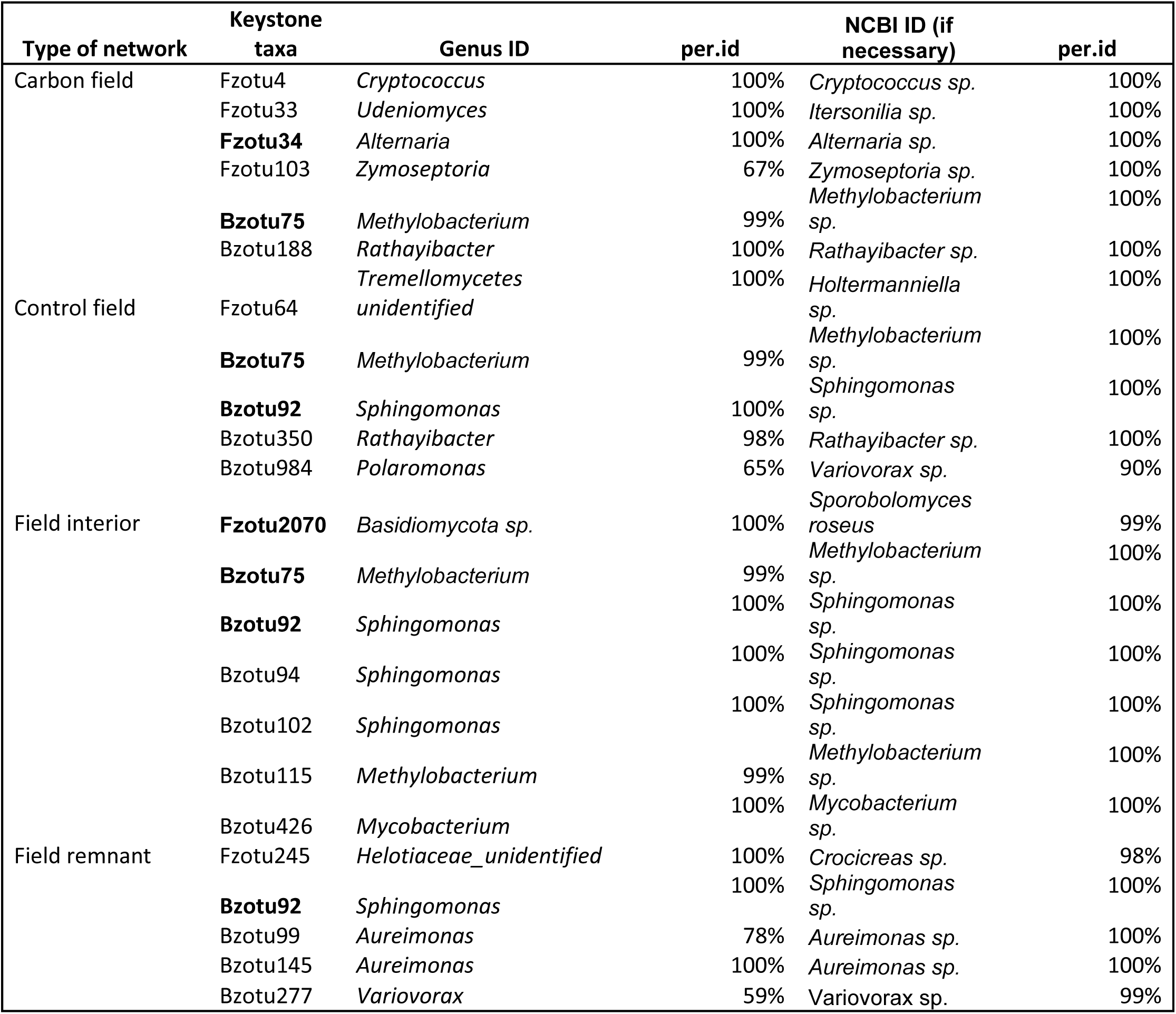
List of the keystone taxa found in the interkingdom networks and their taxonomic identification based on rdp (UNITE and Silva) and NCBI nucleotide BLAST. ZOTUs in bold are keystone taxa shared between networks be them fungal, bacterial or interkingdom. Per.id score indicates the certainty of the taxonomic attribution.

At the fungal-only networks remained relatively well connected under all conditions and exhibited distinct condition specific keystone taxa. Under carbon farming, modules centered on *Neosetophoma* (Fzotu121) and *Zymoseptoria* (Fzotu103; Table 6) whereas in the control network, the main modules were organized around *Cryptococcus* (Fzotu136), *Holtermanniella* (Fzotu64), and *S. roseus* (Fzotu2070). In the field interiors, keystone taxa were *Holtermanniella* (Fzotu64), *Tilletiopsis* (Fzotu143), and *S. roseus* (Fzotu2070) and in the remnant, an unknown taxon (Fzotu147) and two *Alternaria* ZOTUs (Fzotu34, Fzotu70; Table 6).

**Table 6.**
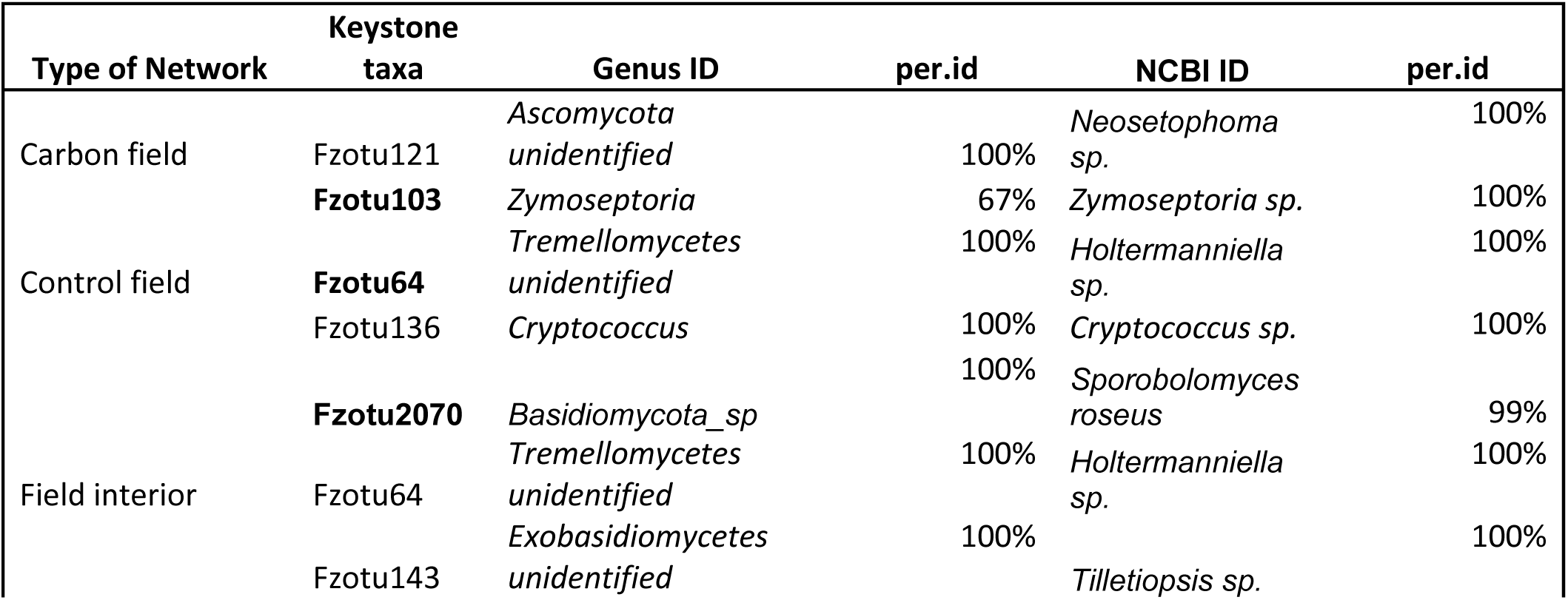
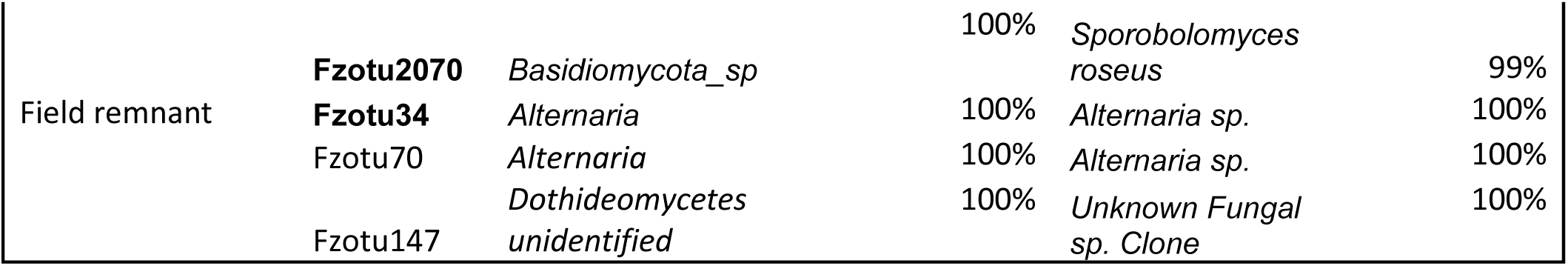
List of the keystone taxa found in the ITS networks and their taxonomic identification based on rdp (UNITE and Silva) and NCBI nucleotide BLAST. ZOTUs in bold are keystone taxa shared between networks be them fungal, bacterial or interkingdom. Per.id score indicates the certainty of the taxonomic attribution.

**Figure 4.**
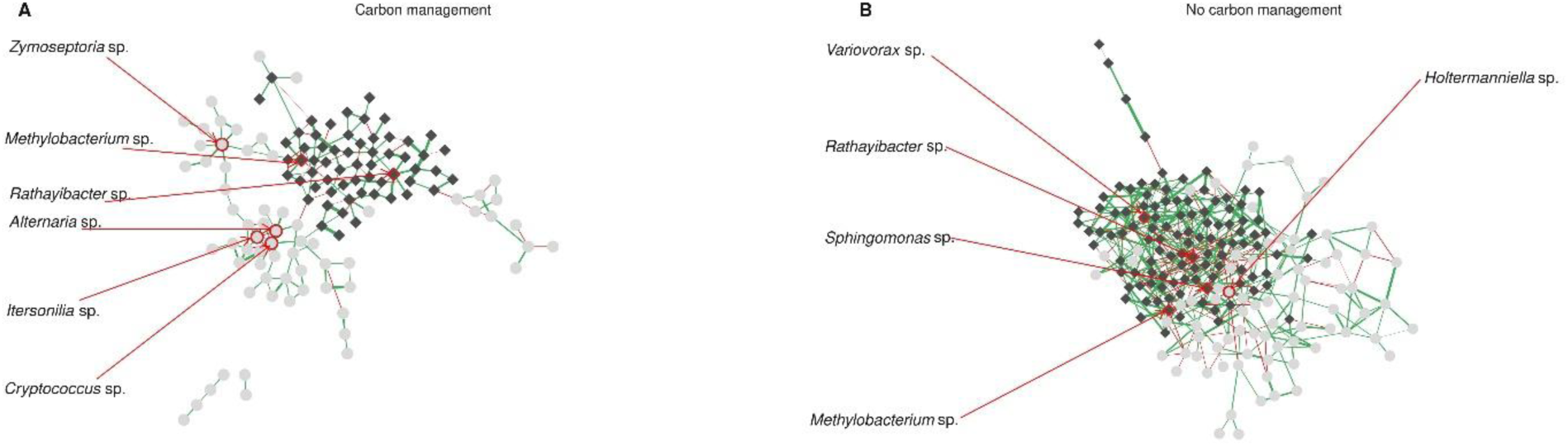
(A) Interkingdom network of co-occurrence and mutual exclusions in the leaf microbiome under carbon management condition and (B) without any carbon management, based on inverse covariance. Nodes in dark grey are bacterial taxa; light grey ones are fungi. The color of the edges indicates positive co-occurrence in green and negative ones in red, the thicker the edges, the stronger the absolute value of inverse covariance coefficient.

By contrast bacterial-only networks were sparse, with no keystone taxa under carbon farming and only single keystone *Methylobacterium* (Bzotu75) under control conditions (Figure S4; Table 7). In the field interior, the bacterial network contained 51 nodes and 48 edges, with *Methylobacterium* (Bzotu75) and *Sphingomonas* (Bzotu68) as keystones. For the remnants, no coherent network was reconstructed; instead, small subunits were observed, with one potential keystone being an unidentified bacterium (Bzotu420; Figure S5; Table 7).

**Table 7.**
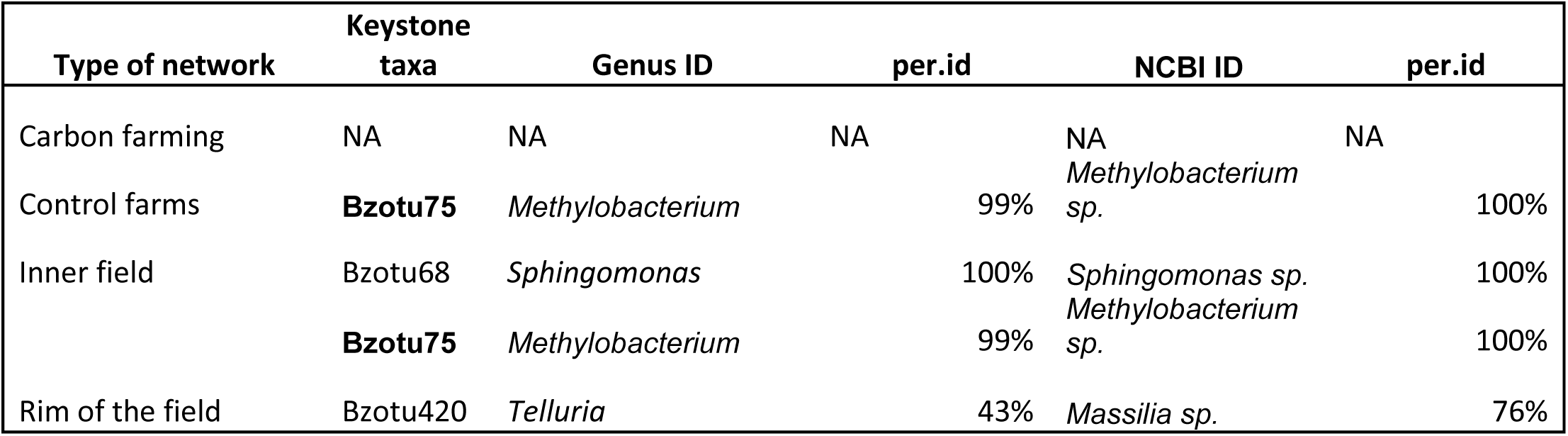
List of the keystone taxa found in the 16S networks and their taxonomic identification based on rdp (UNITE and Silva) and NCBI nucleotide BLAST. ZOTUs in bold are keystone taxa shared between networks be them fungal, bacterial or interkingdom. Per.id score indicates the certainty of the taxonomic attribution.

### 3. Carbon farming and its impact on leaf disease

#### a. Carbon farming practices do not explain the disease load in plant leaves

We examined the influence of carbon farming practices and sampling location (field interior or remnant vegetation) disease symptoms in crops using linear mixed models. Across all disease symptom categories (Supplementary Table S1), the models explained only a small proportion of the observed variation. For example, models for rust (R^2^ = 0.018) and virus symptoms (R^2^ = 0.01506) showed no meaningful effects of carbon farming treatment or sampling location. Sampling location significantly affected rust symptoms (p = 0.0328), although the effect size was small.

Similarly, while a few individual treatments contrasts were significant (e.g., **C**ontrol all-in for Mildew, p = 0.00166), the overall explanatory power of the mildew remained low (R^2^ = 0.04068). No significant treatment effects were detected for leafspot or the remaining disease symptoms. Overall, carbon farming treatment and sampling location explained little of the observed variation in disease symptoms.

#### b. Indicator taxa associated with disease symptoms

Indicator species analysis identified multiple fungal taxa associated with specific disease and herbivory symptoms (Table S4). Several of these corresponded to known or putative plant pathogens (marked with an asterisk), including Pleosporaceae sp. (Zotu12), *Phoma brasiliensis* (Zotu159), *Alternaria eichhorniae* (Zotu78), *A. metachromatica* (Zotu34), *Cladosporium* sp. Chiang 1588 (Zotu80), *Phaeosphaeria* sp. (Zotu69), and *Oculimacula yallundae* (Zotu83).

Rust, mildew, and virus symptoms shared several indicator taxa. Rust symptoms were associated with both pathogenic fungi (e.g., *P. brasiliensis*) and yeasts such as *Cryptococcus* sp., whereas mildew symptoms were characterized by *Cryptococcus* sp., an unidentified Ascomycota, and *P. brasiliensis.* Viral symptoms were associated with *A. eichhorniae* and *Cryptococcus victoriae*.

Herbivore-associated communities also included both pathogenic fungi and yeasts. Large-hole and small-hole damage were associated with *A. metachromatica*, while aphid-infested leaves were associated with *Cladosporium* sp. and *Dissoconium proteae* and leaf miners were characterized with *Phaeosphaeria* sp., *P. brasiliensis*, and yeasts such as *Dioszegia hungarica*. Gall symptoms exhibited the greatest diversity of fungal indicator taxa including *Stagonospora pseudovitensis*, *O. yallundae*, *Pleosporales* sp. and several yeast taxa.

Bacterial indicator taxa also varied among symptom types (Supplementary Table S5). Rust symptoms were associated with the greatest diversity of bacterial taxa, dominated by several *Sphingomonas*, *Methylobacterium*, *Hymenobacter* ZOTUs, together with *Alpinimonas*, *Frondihabitans*, *Curtobacterium*, and *Microterricola*. Mildew symptoms likewise showed strong association with *Sphingomonas*, represented by at least five distinct ZOTUs, together with *Aureimonas*. Other symptom categories were characterized by fewer indicator taxa, including *Clavibacter* for large-hole damage, Pantoea for thrips damage, Massilia for scraping damage, and Sphingomonas together with *Hephaestia* for mite-associated communities.

##### Partial least squares path modeling of environmental, microbial, and disease interactions

To disentangle the direct and indirect relationships among environmental factors, microbial communities, and disease load, we applied partial least squares path modeling (PLS-PM; Figure 5). The PLS-PM analysis revealed direct and indirect relationships among environmental variables, microbial community composition, and disease load (Figure 5). Farm identity had the strongest effects in the model, showing a negative association with fungal communities (β = −0.50) and a positive association with bacterial communities (β = 0.40). Carbon farming treatment showed weak negative associations with both fungal (β = −0.13) and bacterial (β = −0.16) communities. Sampling location (field interior or remnant vegetation) was only weakly associated with fungal communities (β = 0.008) but shoved a moderate positive association with bacterial communities (β = 0.30).

**Figure 5.**
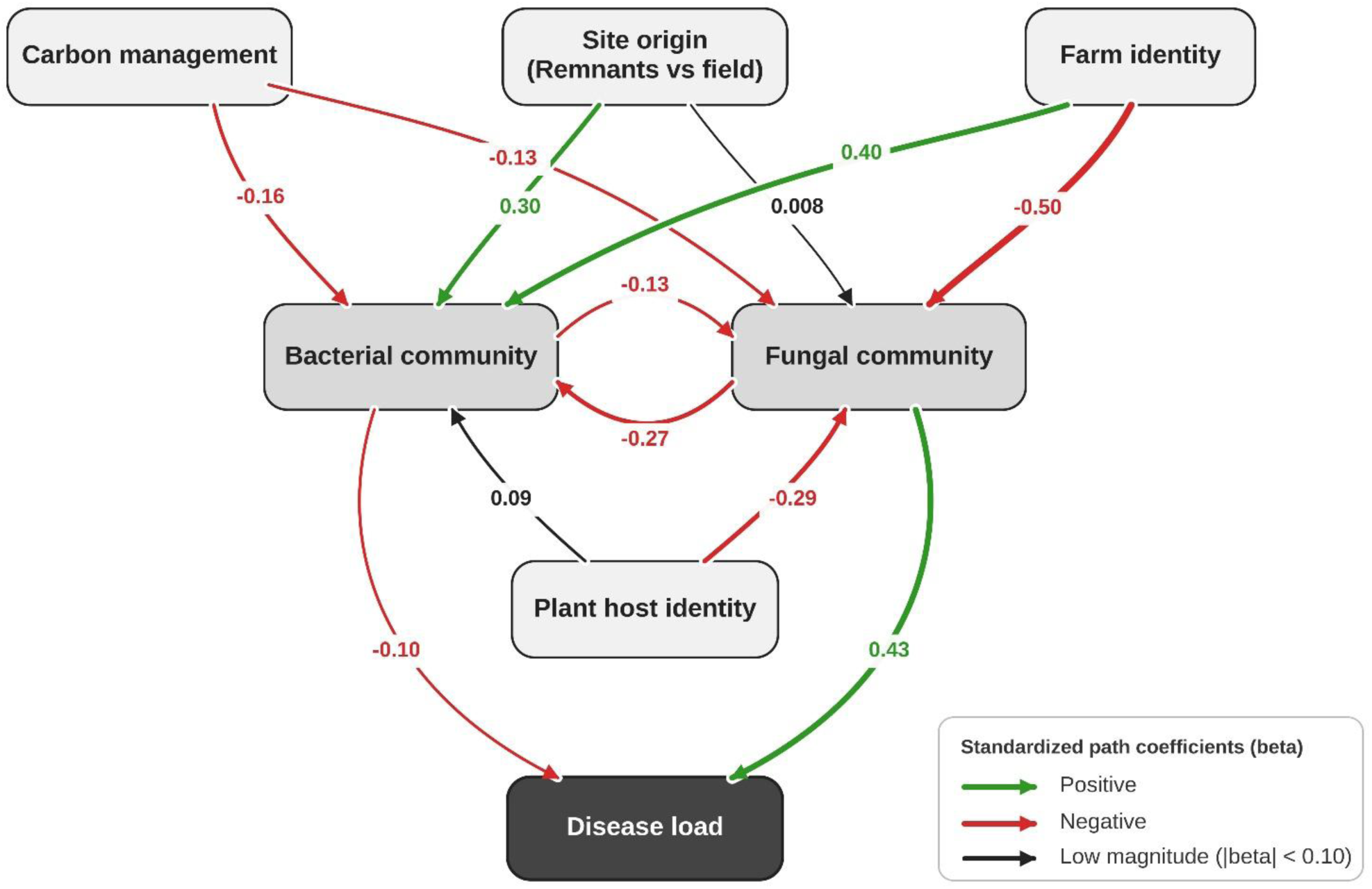
Partial least squares path modeling (PLS-PM) of our experimental system illustrating the relationships among carbon farming, sampling location (field interior or remnant vegetation), host plant identity, farm identity, bacterial and fungal leaf community composition and disease load. Arrows indicate hypothesized causal paths, with numbers showing standardized path coefficient (β). Arrow thickness is proportional to the absolute value of the coefficient with a color code; red for negative coefficient and green for positive one, black is for low values of β.

Host plant identity was negatively associated with fungal communities (β = −0.29) and weakly positively associated with bacterial communities (β = 0.09). The relationship between fungal and bacterial communities was negative in both directions, with fungi negatively associated with bacteria (β = −0.27) and bacteria negatively associated with fungi (β = −0.13). Disease load was positively associated with fungal communities (β = 0.43), but only weakly negatively associated with bacterial communities (β = −0.10).

Overall, the PLS-PMs analysis suggests that farm identity was the strongest predictor of microbial community composition, while host plant identity and carbon farming treatment made smaller contributions. Fungal communities were more strongly associated with disease loads than bacterial communities, the latter appearing to influence disease primarily via their interactions with fungi.

## Discussion

In this study, we investigated whether carbon farming practices alter the leaf microbiome and plant health across agricultural systems. We found that carbon farming practices exert significant effects on both fungal and bacterial communities inhabiting phyllosphere communities, with fungal assemblages showing stronger and more consistent responses (Figure 2). Despite these management-driven shifts, host identity emerged as the dominant factor structuring the microbiome, underscoring the strong influence of plant species on microbial composition. Interestingly, control plots—those not subjected to carbon amendments— generally supported more indicator taxa and denser microbial co-occurrence networks than carbon farming treatments. Meanwhile, disease load was only weakly associated with carbon farming or sampling location indicating that additional environmental or biological drivers likely govern disease dynamics. Collectively, these findings suggest that carbon farming practices are associated with shifts in aboveground microbial assemblages, highlighting how management practices designed to enhance soil carbon sequestration may also influence aboveground microbial communities.

### 1. The influence of carbon farming on the microbial diversity and structure of plant phyllosphere

Our results show that carbon farming practices significantly altered both bacterial and fungal diversity in the phyllosphere, with fungal communities responding more strongly and consistently than bacterial communities. This suggests that management practices designed to enhance soil carbon sequestration influence the plant–microbe continuum, potentially by modifying nutrient availability, root exudation profiles, or plant physiological traits that in turn affect leaf-surface colonization. Similar cross-compartment effects have been observed when soil amendments or compost additions reshaped aboveground microbial communities through altered carbon fluxes and plant signaling (Kelly et al. 2022; Heinemann et al. 2023). The strong host identity effect observed here aligns with previous evidence that plant traits and genotypes filter microbial assemblages (Patel et al. 2015; Bashir et al. 2022). However, the magnitude of this filtering varied across carbon treatments, indicating that the outcome of management interventions could depend on host-specific physiological responses. For example, species with higher leaf nutrient turnover or cuticular permeability may transmit soil-derived carbon or microbial signals more efficiently, resulting in stronger phyllosphere shifts (Kelly et al. 2022; Heinemann et al. 2023).

Adaptive grazing produced some of the strongest shifts in phyllosphere community composition, likely because grazing introduces repeated physical disturbance to plant tissues. Leaf damage caused by grazing creates entry points for microorganisms, alters plant physiology, and increases the abundance of senescing tissue, all of which may favor necrotrophic fungi and opportunistic pathogens observed *P. phragmiticola*, *Pleosporales sp*., *Oculimacula sp*., *Blumeria graminis* and *Kabatiella bupleuri* (Table S2) (Carson 2005; Bills, Menéndez, and Platas 2012; Ramanauskienė et al. 2014). The other carbon farming practice that displayed major differences in fungal community structure when compared to others was the ley mixture. Ley mixtures likely supported distinct fungal communities because multiple host species co-occurred within the same field, allowing the community to integrate the host-specific microbial assemblages associated with different plant species (Yao et al. 2019; M. Li et al. 2022; Köhler et al. 2025). Furthermore, mixed leys combine plants with contrasting leaf surface traits known to affect microbial communities, such as surface pH, wax chemistry, trichome density and cuticle thickness (Aragón, Reina-Pinto, and Serrano 2017; Kusstatscher et al. 2020; Lopez-Gonzalez et al. 2025). The fact that ley mixture was also consistently associated with yeasts, saprotrophs and mild pathogens (Table S2) is coherent with what we could expect from dense, multi-species canopies that can produce stable humidity gradients and variable microclimates (Sentelhas et al. 2008; Gouka, Raaijmakers, and Cordovez 2022; He et al. 2025). Interestingly, the two carbon farming practices that differed most from the remaining treatments - adaptive grazing and ley mixture- also shared two ecological characteristics: recurrent disturbances and perennial root systems. Together, these features may promote continual microbial recolonization from soil and neighboring vegetation, maintaining a dynamic and spatially heterogeneous phyllosphere microbiome.

### 2. Contrasting ecological roles of *Alternaria sp.* and *Sporobolomyces roseus*

Network analysis revealed consistent keystone Zotus shared between different conditions and between interkingdom and intra-kingdom dynamics (Figure S5; Tables 5, 6 and 7). Two taxa were repeatedly identified as keystone taxa across interkingdom and within-kingdom networks: *Sprobolomyces roseus* (FZotu 2070) and *Alternaria sp.* (FZotu 34). Their repeated occurrence suggests that they may play important ecological roles within phyllosphere microbial communities.

*S. roseus* is a species complex of psychrotrophic yeasts (Rusinova-Videva et al. 2024; Białkowska et al. 2018), frequently described in the wheat phyllosphere since 1976 (Bashi and Fokkema 1976; Fokkema and Meulen 1976; Bashi and Fokkema 1977). However subsequent studies found it in association of a wide range of plants, always in their phyllosphere, like in other crops (rice, maize) and grasses, but also trees and herbs such as *Populus*, *Acer* and *Artemisia* (Bai 2002; Q.-M. Wang and Bai 2004), indicating that the yeasts from the *S.roseus* complex to be ubiquitous with no particular host specificity. In our study, *S. roseus* is found as a keystone species in the Field interior at interkingdom level and control farms when considering only the fungal community. This is consistent with previous reports because the sampled crop communities consisted predominantly of *Poaceae*. In the leaf system, *S. roseus* is known to be an antagonist to several other species of pathogenic fungi. It reduces infection of specific fungal pathogens such as *Septoria nodorum*, and *Cochliobolus sativus* (Fokkema and Meulen 1976; Bashi and Fokkema 1977), and overall tends to decrease leaf necrosis in wheat (Fokkema and De Nooij 1981). More recent studies focus on phyllosphere yeasts broadly, and species-specific evidence remains limited (Gouka, Raaijmakers, and Cordovez 2022; Rangel and Leveau 2024). One possible explanation for this pattern is that *S. roseus* suppresses fungal pathogens either directly or indirectly by promoting stable yeast biofilms that occupy phyllosphere niches before pathogen establishment. (Dickinson and O’Donnell 1977; Frossard, Fokkema, and Tietema 1983; Sapkota, Jørgensen, and Nicolaisen 2017).

*Alternaria spp*. are filamentous fungal species that occurs as both saprotrophs and opportunistic pathogens (Dai et al. 2022; Fernandes, Casadevall, and Gonçalves 2023). Our results identified *Alternaria sp*. as a recurrent keystone taxon under carbon farming treatments at interkingdom and intra-kingdom level. Its abundance also appears to increase in other studies under disturbance in the leaf system, and leaf wetness (Vloutoglou and Kalogerakis 2000; Pleysier et al. 2006). *Alternaria* is often described as a genus of phyllospheric fungi that occurs in the late successional stage of the leaf microbial colonization, usually replacing early colonizing yeast, benefiting from carbon made available from cell death (Voříšková and Baldrian 2013; Dai et al. 2022). *Alternaria spp*. also displays some facilitation for a cohort of saprophytic and opportunistic fungi in the leaf system like *Cladosoprium*, *Epicocum* and *Phoma* that tends to show higher relative abundance on leaves already infected by *Alternaria* (Tao et al. 2021; Dai et al. 2022). Taken together, these observations suggest that *Alternaria spp*. and *S. roseus* represent two contrasting ecological states of the phyllosphere. Whereas *Alternaria* was associated with carbon farming treatments and remnant vegetation—conditions likely characterized by greater disturbance and tissue turnover— *S. roseus* was primarily associated with field communities and control plots, consistent with previous reports linking this yeast to stable cereal phyllospheres. Rather than directly antagonizing one another, these taxa may indicate alternative microbial assemblages associated with contrasting environmental conditions. It is interesting to note that *Alternaria spp*. was also flagged as a keystone taxon in the network made from the field remnants and not the one from the Field interior. This observation is consistent with the hypothesis that remnant vegetation may function as reservoirs of microbial diversity and contribute to the spatial structuring of phyllosphere communities (Haas et al. 2011; Keesing and Ostfeld 2021; Hasanaliyeva et al. 2024).

### 3. Bacterial dynamics and the importance of *Methylobacterium sp*. in the phyllosphere

Interkingdom co-occurrence networks revealed a clear separation between fungal and bacterial communities with relatively few cross-kingdom associations, particularly under carbon farming treatments. This contrasts with previous studies about plant phyllospheric interactome, as most of them underlined intricacy between fungal and bacterial dynamics(Agler et al. 2016; Bowsher et al. 2021; Zhao et al. 2023). Particularly when considering that the labile interface of fungal hyphae can be serve as “bacterial highways” to move more easily in their environment (Warmink et al. 2011; Simon et al. 2015; Ruan et al. 2022). Because co-occurrence networks are sensitive to differences in statistical methodology and network inference algorithms, comparisons among studies should be interpreted cautiously (Kurtz et al. 2015; Gloor and Reid 2016). That said, the co-occurrence patterns observed in our study between fungi and bacteria support the possibility that the two kingdoms react very differently to perturbation and environmental variations as described by (Liu et al. 2024) in a forest environment.

Among the bacterial taxa, *Methylobacterium sp*. emerged as the most consistent keystone, occupying central positions in both bacterial and interkingdom networks. Members of the genus Methylobacterium are widespread components of the plant microbiome, occurring on leaves, roots, and seeds (Palberg et al. 2022; Sanjenbam, Shivaprasad, and Agashe 2022; Xiong et al. 2024; Grossi et al. 2025. Species of *Methylobacterium* tend to be diazotrophs and acts as key symbionts in the plant leaf microbiome as they are often linked to significant increase in crop yield (Zhang et al. 2021; Sanjenbam, Shivaprasad, and Agashe 2022) or offers protection against pathogenic attacks (Innerebner, Knief, and Vorholt 2011). The only interkingdom association involving *Methylobacterium* (BZotu75) connected it with *Ramularia hydrangeae*, a known foliar pathogen (Park and Shin 2016; Bakhshi, Zare, and Druzhinina 2025). Although co-occurrence does not imply direct interaction, this association may reflect shared occupancy of the leaf surface rather than ecological dependence. Similar co-occurrence has previously been reported across diverse wild flowering plant species (Ramakrishnan et al. 2024). Because our analyses are based on co-occurrence networks the ecological role of *Methylobacterium* remains speculative. Nevertheless, its repeated identification as a keystone taxon in both bacterial and interkingdom level, together with its documented beneficial effects on plants, highlights this genus as a promising target for future studies of phyllosphere microbiomes in agricultural systems.

## Conclusion

Our study shows that soil carbon farming practices influence aboveground microbial communities, with fungal communities responding more strongly than bacterial communities. Yet, plant host identity remained the dominant factor shaping phyllosphere composition. Control plots supported more indicator taxa and denser microbial co-occurrence networks than carbon farming treatments, suggesting that carbon farming practices may alter microbial network complexity. Disturbance-based practices such as adaptive grazing and ley mixtures generated the most distinct microbial communities, suggesting that management practices designed to enhance soil carbon sequestration can influence leaf microbial assembly. Network analyses repeatedly identified *Alternaria* spp. and *Sporobolomyces roseus* as contrasting keystone taxa associated with different phyllosphere community structures. *Methylobacterium* spp. also emerged as an important bacterial structuring taxa across treatments. Overall, these findings show that carbon farming practices influence phyllosphere microbial communities, altering community composition, network structure, and the distribution of potential keystone taxa. Future work should investigate the mechanisms by which carbon farming practices influence leaf microbiome assembly and determine how these changes affect plant health and the long-term sustainability of agricultural systems.

## Acknowledgements

We’re grateful to the Baltic Sea Action Group and Tuomas Mattila for assisting in establishing the farm collaboration network. We thank Krista Raveala, Anna Välkki, and Laura Kares for assisting in the field work, and Ilona Peltoniemi for assisting in the laboratory. We thank Funding for this project was provided by the Research Council of Finland (STN MULTA; 327222) to A-L.L., The Finnish Innovation Fund Sitra to A.-L.L. and HS and European Research Council (AdG 101097545 Co-EvoChange to A.-L.L.).

