## Supplementary figures and images for "Field Plant Biodiversity and Carbon Farming Shape Leaf Microbiomes"

### Figure S1

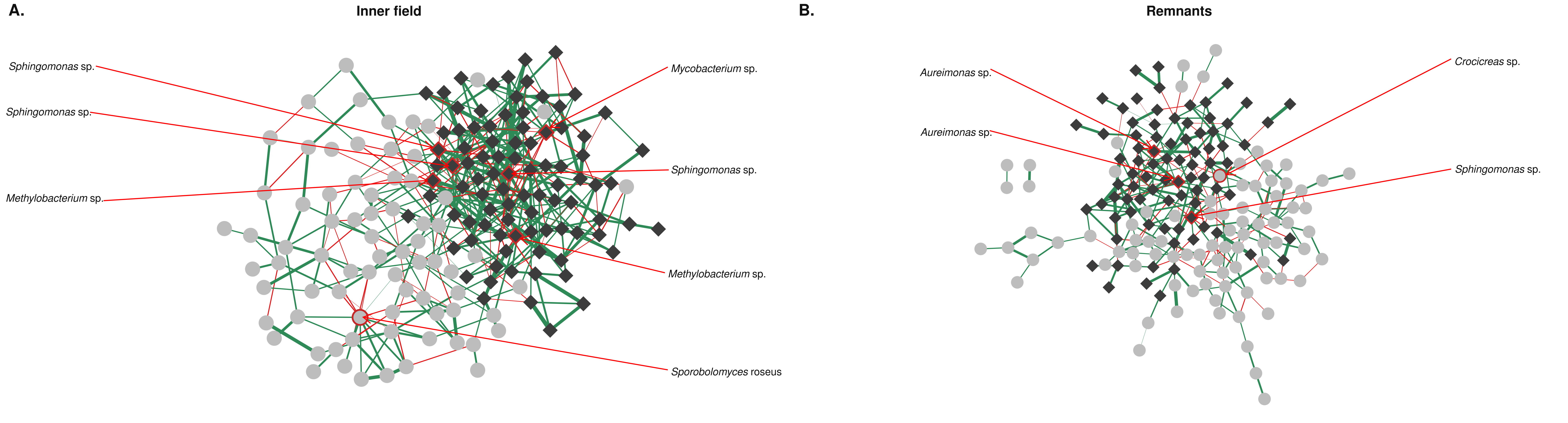

### Figure S2

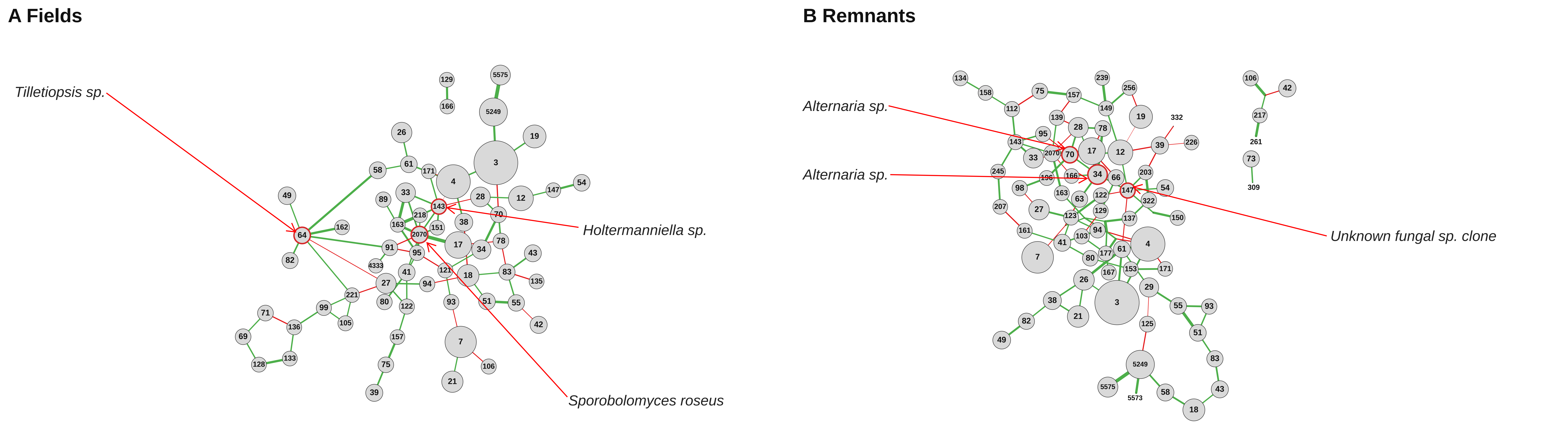

### Figure S3

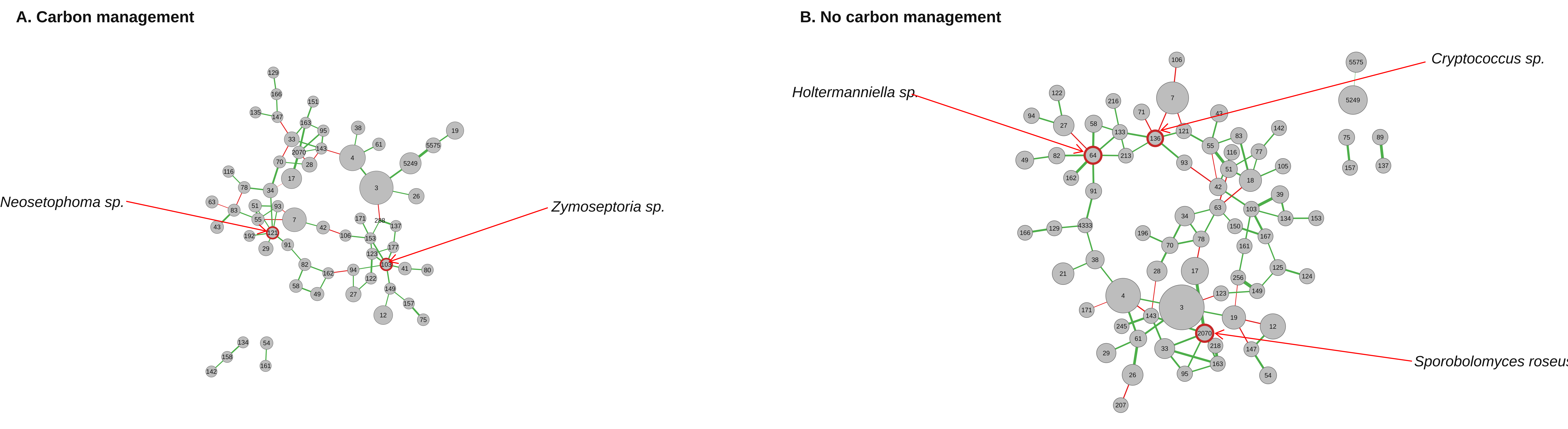

### Figure S4

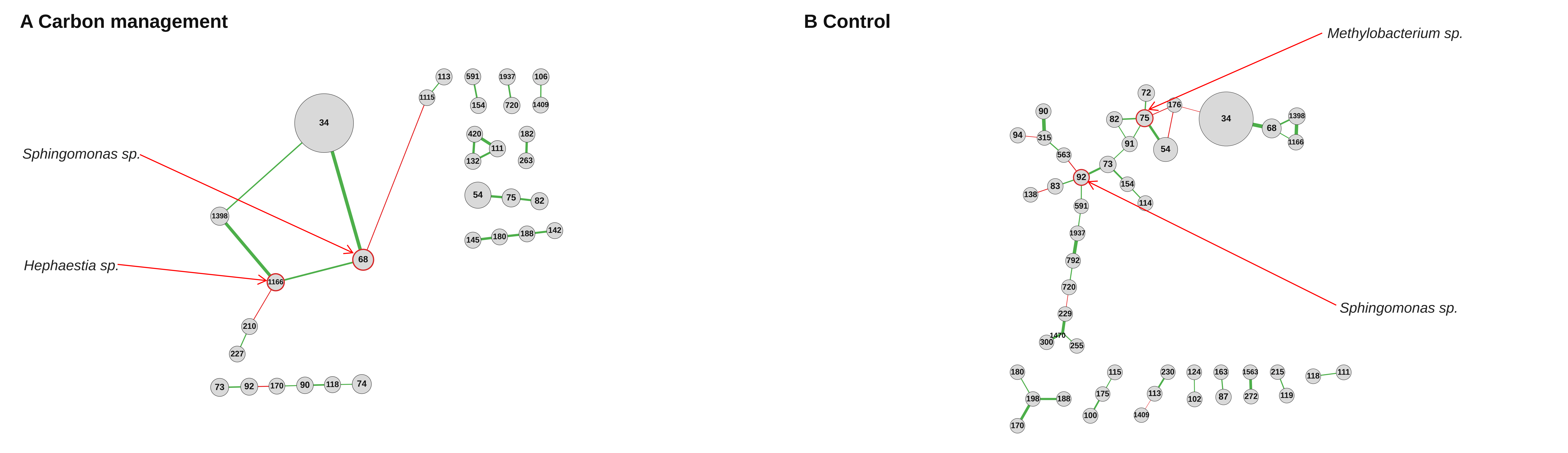

### Figure S5

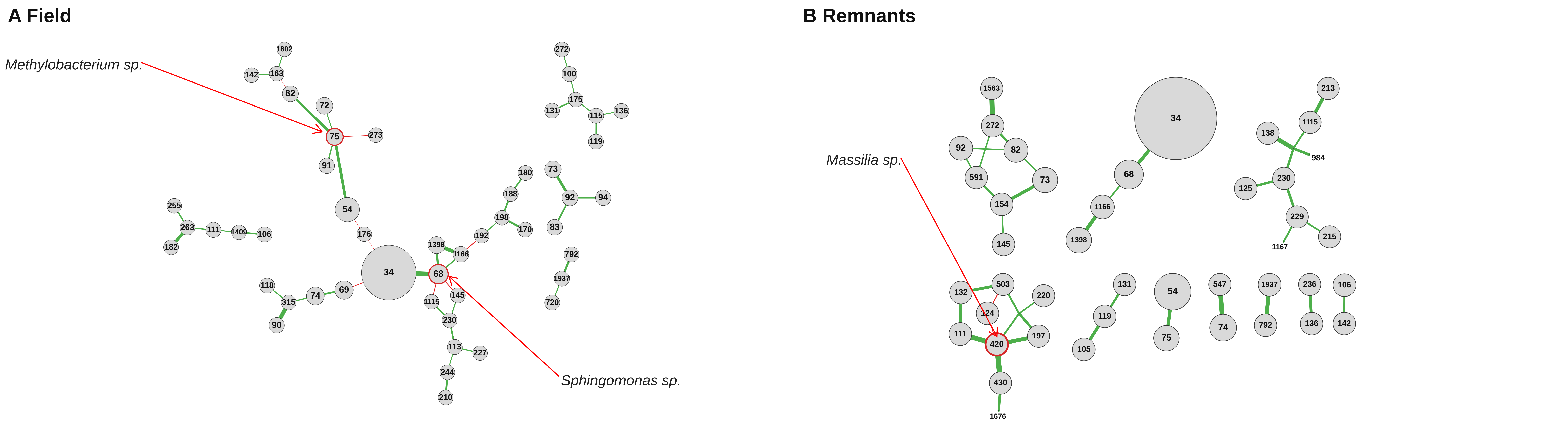

### Figure S6

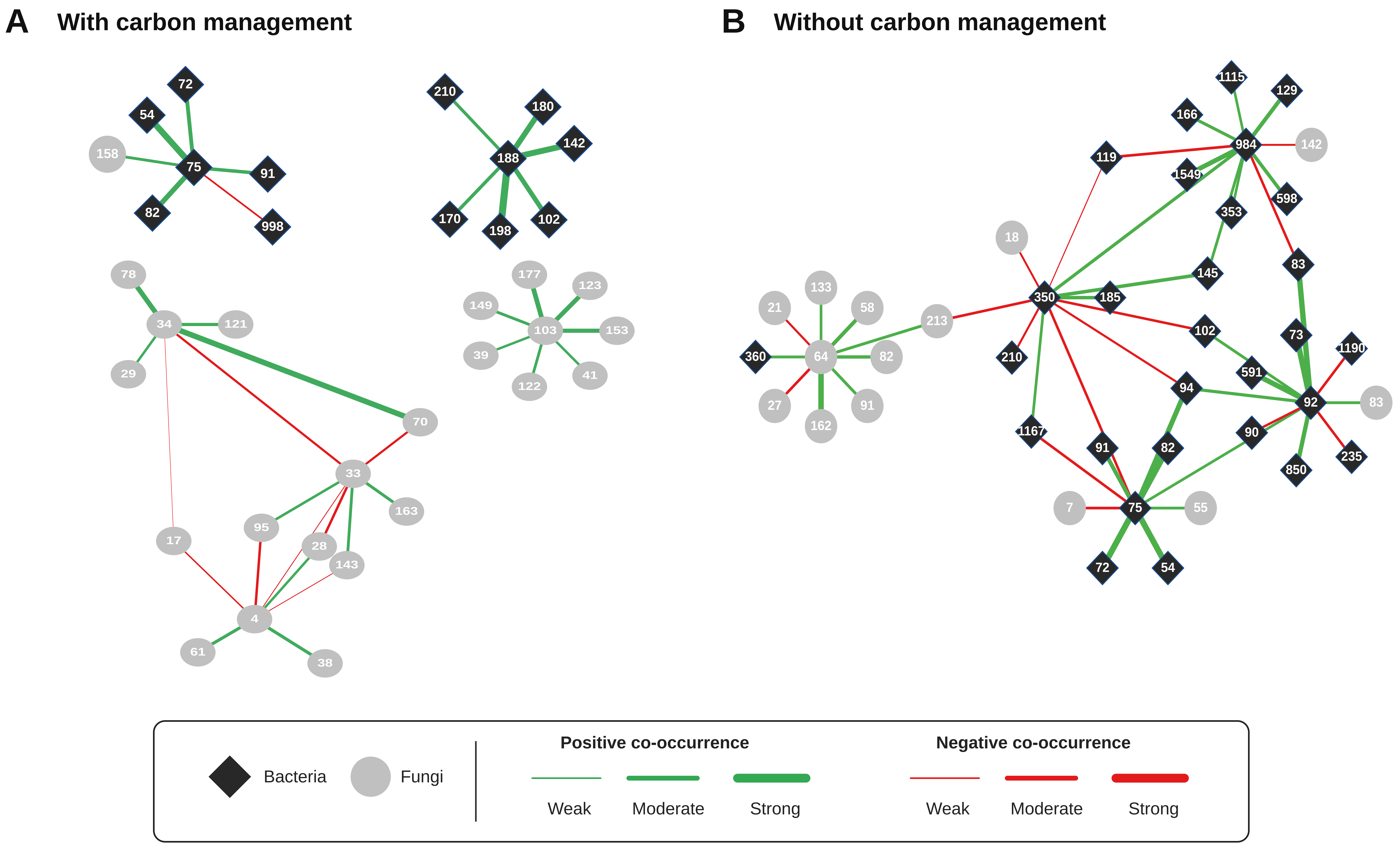
