## Supplementary Tables for "Field Plant Biodiversity and Carbon Farming Shape Leaf Microbiomes"

| **Symptom** | **Description** | **Causal Agents (Kingdom)** | **Representative Species** | **Reference** |
| --- | --- | --- | --- | --- |
| **Rust** | Red/orange pustules on leaves, stems, or other plant parts | Fungi | *Puccinia graminis (wheat stem rust), Puccinia triticina (wheat leaf rust), Uromyces appendiculatus (common bean rust)* | Wanyera et al. (2006); Kolmer et al. (2009) |
| **Mildew** | White, powdery patches on leaves, stems, and flowers | Fungi | *Erysiphe spp. (powdery mildew on cereals and legumes), Blumeria graminis (powdery mildew on grasses, wheat, and barley)* | Dean et al. (2012); Glawe (2008) |
| **Virus** | Discoloration of leaves and plant tissues | Viruses | Barley yellow dwarf virus (BYDV), Potato virus Y (PVY), Potato mop-top virus (PMTV) | D'Arcy (1995); Jones (2004) |
| **Leaf Spot** | Discrete circular or irregular lesions in the leaves that vary in color from black, brown, or gray to red and yellow | Bacteria, Fungi, Viruses | *Xanthomonas spp., Pseudomonas syringae, Erwinia spp.; Alternaria spp. (brassicas), Cercospora spp. (legumes), Septoria spp. (brassicas); Pyrenophora teres (barley), Cochliobolus sativus (barley, oat), Rhynchosporium secalis (barley), Stagonospora nodorum (wheat), Pyrenophora tritici-repentis (wheat), Pyrenophora chaetomioides (oat); Tomato spotted wilt virus (TSWV), Cucumber mosaic virus (CMV)* | Agrios (2005); Windels et al. (1998); Jalli et al. (2011) |
| **Large Hole** | Recognized as shot-hole patterns and from slime marks left by insects | Insects (caterpillars, beetles, grasshoppers), Mollusks (slugs, snails), Bacteria, Fungi | *Xanthomonas spp., Coryneum spp., Alternaria solani* | Agrios (2005); Jain et al. (2019) |
| **Small Hole** | Same identification steps as with large holes | Insects (flea beetles, caterpillars, leaf miners, thrips, weevils), Bacteria, Fungi | *Xanthomonas spp., Coryneum spp.; leaf spots can become holes after infection by Alternaria, Cercospora, or Septoria spp.* | Agrios (2005); Capinera (2001) |
| **Window Feeding** | The top layer of the leaf is eaten leaving the lower layer intact, creating a windowpane appearance | Insects (leaf miners, caterpillars, slugs, snails, thrips, beetles), Bacteria, Fungi | Fungal and bacterial leaf spots can become windows | Capinera (2001); Agrios (2005) |
| **Thrips Damage** | Silvery discoloration, stippling, tiny black spots of frass (insect excrement) | Insects (thrips), Bacteria, Fungi | — | Lewis (1997); Agrios (2005) |
| **Aphid Damage** | Yellowing and wilting of plants, accumulation of sticky honeydew secreted by aphids, presence of mosaic viruses transmitted by aphids | Insects (aphids), Bacteria, Fungi, Viruses | Black mold and mosaic viruses, tospoviruses | Blackman & Eastop (2000); Agrios (2005) |
| **Mining** | Light-colored mines or tunnels formed by miners | Insects (leaf miners), Bacteria, Fungi | — | Capinera (2001); Agrios (2005) |
| **Scraping** | Whitish, translucent or silvery patch | Insects (thrips, spider mites, beetle larvae), Mollusks (slugs, snails), Bacteria, Fungi, Mechanical injury | Powdery mildew pathogens | Agrios (2005); Capinera (2001) |
| **Moth Damage** | Root damage, mines created by larvae, silk production | Insects (moths - Lepidoptera), Bacteria, Fungi | — | Capinera (2001); Agrios (2005) |
| **Spittlebug Foam** | Bubbly white foam covering the insect and plant tissues | Insects (spittlebug - froghopper), Bacteria, Fungi | — | Hamilton & Morales (1992); Agrios (2005) |
| **Mite Damage** | Yellow/white stipples in leaves, webbing (spider mites), leaf distortion, stunted growth | Arachnids (mites), Bacteria, Fungi | Spider mites (Tetranychidae), broad mites (Polyphagotarsonemus latus), russet mites (Aculops spp.) | Jeppson et al. (1975); Agrios (2005) |
| **Galls** | Deformed growths on the plants, discoloration to yellowish when caused by bacteria/fungi, stunted growth | Insects, Mites, Fungi, Bacteria, Nematodes | Bacteria: Agrobacterium tumefaciens; Fungi: Ustilago spp. (grasses) | Agrios (2005); Harris & Pitzschke (2015) |
| **Caterpillar Damage** | Chewed leaves, frass (excrements), silken webbing, visible caterpillars | Insects (caterpillars), Bacteria, Fungi | Leaf-mining caterpillars (various species) | Capinera (2001); Agrios (2005) |
| **Other Leaf Symptoms** | Yellowing (chlorosis), curling or distortion, brown or blackened leaf tips, scorched appearance, leaf spots, holes or tearing | Various pathogens and pests as listed above | — | Agrios (2005) |

**Table S1.** Classification of the disease symptoms in our study according to the literature.

| **Group** | **ZOTU** | **Genus** | ***species*** | **Accession** | **Genus Confidence (%)** |
| --- | --- | --- | --- | --- | --- |
| adaptive grazing | Zotu134 | Ascomycota unidentified | *Ascomycota sp.* | SH216151.06FU | 100 |
| adaptive grazing | Zotu75 | Phaeosphaeriaceae unidentified | *Phaeosp.haeriaceae sp.* | SH206980.06FU | 100 |
| adaptive grazing | Zotu453 | Phaeosphaeriaceae unidentified | *Phaeosp.haeriaceae sp.* | SH206980.06FU | 100 |
| adaptive grazing | Zotu327 | Vibrisseaceae unidentified | *Vibrisseaceae sp.* | SH234171.06FU | 36 |
| adaptive grazing | Zotu263 | Kabatiella | *Kabatiella bupleuri* | SH229248.06FU | 62 |
| adaptive grazing | Zotu451 | Cistella | *Cistella sp.* | SH209363.06FU | 98 |
| adaptive grazing | Zotu39 | Ascomycota unidentified | *Ascomycota sp.* | SH216151.06FU | 100 |
| adaptive grazing | Zotu337 | Phaeosphaeria | *Phaeosp.haeria sp.* | SH199976.06FU | 55 |
| adaptive grazing | Zotu382 | Phialocephala | *Phialocephala fluminis* | SH209755.06FU | 20 |
| adaptive grazing | Zotu331 | Rhexocercosporidium | *Rhexocercosp.oridium panacis* | SH204719.06FU | 79 |
| adaptive grazing | Zotu157 | Phaeosphaeria | *Phaeosp.haeria phragmiticola* | SH199979.06FU | 49 |
| adaptive grazing | Zotu92 | Blumeria | *Blumeria graminis* | SH195226.06FU | 100 |
| adaptive grazing | Zotu40 | Cryptococcus | *Cryptococcus sp.* | SH223310.06FU | 100 |
| adaptive grazing | Zotu106 | Pleosporales unidentified | *Pleosp.orales sp.* | SH233950.06FU | 100 |
| adaptive grazing | Zotu234 | Trichopeziza | *Trichopeziza mollissima* | SH239999.06FU | 99 |
| adaptive grazing | Zotu284 | Ascomycota unidentified | *Ascomycota sp.* | SH216151.06FU | 58 |
| adaptive grazing | Zotu328 | Tremellomycetes unidentified | *Tremellomycetes sp.* | SH203518.06FU | 100 |
| adaptive grazing | Zotu179 | Oculimacula | *Oculimacula sp.* | SH204743.06FU | 91 |
| all-in | Zotu690 | Pleosporales unidentified | *Pleosp.orales sp.* | SH227424.06FU | 97 |
| all-in | Zotu316 | Dissoconium | *Dissoconium proteae* | SH197939.06FU | 100 |
| all-in | Zotu409 | Venturia | *Venturia tremulae* | SH207401.06FU | 99 |
| Control adaptive grazing | Zotu255 | Sarocladium | *Sarocladium sp.* | SH191583.06FU | 100 |
| Control adaptive grazing | Zotu269 | Myrothecium | *Myrothecium gramineum* | SH199496.06FU | 85 |
| Control adaptive grazing | Zotu246 | Phoma | *Phoma pasp.ali* | SH233949.06FU | 100 |
| Control adaptive grazing | Zotu18 | Incertae sedis unidentified | *Pleosp.orales sp.* | SH233951.06FU | 100 |
| Control adaptive grazing | Zotu297 | Stagonospora | *Stagonosp.ora pseudovitensis* | SH199974.06FU | 50 |
| Control adaptive grazing | Zotu189 | Acremonium | *Acremonium sp.* | SH222251.06FU | 64 |
| Control adaptive grazing | Zotu204 | Sordariomycetes unidentified | *Sordariomycetes sp.* | SH219629.06FU | 100 |
| Control adaptive grazing | Zotu184 | Hymenula | *Hymenula cerealis* | SH204721.06FU | 99 |
| Control adaptive grazing | Zotu564 | Myrothecium | *Myrothecium sp.* | SH199487.06FU | 100 |
| Control adaptive grazing | Zotu267 | Hyaloscyphaceae unidentified | *Hyaloscyphaceae sp.* | SH197899.06FU | 31 |
| Control adaptive grazing | Zotu108 | Ascomycota unidentified | *Ascomycota sp.* | SH241083.06FU | 82 |
| Control adaptive grazing | Zotu61 | Cryptococcus | *Cryptococcus sp.* | SH198056.06FU | 100 |
| Control adaptive grazing | Zotu182 | Cryptococcus | *Cryptococcus aff amylolyticus* | SH223364.06FU | 100 |
| Control all-in | Zotu82 | Cryptococcus | *Cryptococcus chernovii* | SH216501.06FU | 100 |
| Control all-in | Zotu725 | Cryptococcus | *Cryptococcus friedmannii* | SH235990.06FU | 100 |
| Control all-in | Zotu2383 | Cryptococcus | *Cryptococcus chernovii* | SH216501.06FU | 100 |
| Control all-in | Zotu4333 | Cryptococcus | *Cryptococcus sp.* | SH198056.06FU | 99 |
| Control all-in | Zotu26 | Cryptococcus | *Cryptococcus victoriae* | SH198055.06FU | 100 |
| Control all-in | Zotu679 | Ascomycota unidentified | *Ascomycota sp.* | SH194662.06FU | 47 |
| Control cover crop | Zotu54 | Ascomycota unidentified | *Ascomycota sp.* | SH241083.06FU | 65 |
| Control leymix | Zotu589 | Phaeosphaeria | *Phaeosp.haeria sp.* | SH227803.06FU | 39 |
| Control leymix | Zotu218 | Basidiomycota unidentified | *Basidiomycota sp.* | SH196706.06FU | 100 |
| Control leymix | Zotu551 | Itersonilia | *Itersonilia perplexans* | SH217649.06FU | 100 |
| Control leymix | Zotu467 | Stagonospora | *Stagonosp.ora pseudovitensis* | SH199974.06FU | 99 |
| Control leymix | Zotu665 | Leptosphaeria | *Leptosp.haeria doliolum* | SH228249.06FU | 99 |
| Control leymix | Zotu553 | Phaeosphaeria | *Phaeosp.haeria caricicola* | SH227808.06FU | 70 |
| Control leymix | Zotu225 | Ascomycota unidentified | *Ascomycota sp.* | SH206984.06FU | 98 |
| Control leymix | Zotu633 | Dioszegia | *Dioszegia hungarica* | SH196961.06FU | 96 |
| Control leymix | Zotu615 | Tremellales unidentified | *Tremellales sp.* | SH198016.06FU | 100 |
| Control leymix | Zotu954 | Udeniomyces | *Udeniomyces pannonicus* | SH217650.06FU | 91 |
| leymix | Zotu143 | Exobasidiomycetes unidentified | *Exobasidiomycetes sp.* | SH230059.06FU | 100 |
| leymix | Zotu95 | Dioszegia | *Dioszegia butyracea* | SH196966.06FU | 100 |
| leymix | Zotu418 | Dioszegia | *Dioszegia butyracea* | SH196966.06FU | 61 |
| leymix | Zotu144 | Pleosporales unidentified | *Pleosp.orales sp.* | SH231238.06FU | 100 |
| leymix | Zotu521 | Ascomycota unidentified | *Ascomycota sp.* | SH225340.06FU | 60 |
| leymix | Zotu53 | Golovinomyces | *Golovinomyces sp.* | SH212551.06FU | 100 |
| leymix | Zotu79 | Helotiaceae unidentified | *Helotiaceae sp.* | SH209325.06FU | 44 |

**Table S2.** Indicator species analysis of the plant leaf fungal communities in function of the carbon farming practices. Indicator Zotus are displayed with their taxonomic attribution according to UNITE. α = 0.05, n = 207.

| **Group** | **ZOTU** | **Genus** | **Genus Confidence (%)** | **stat** | **p.value** |
| --- | --- | --- | --- | --- | --- |
| adaptive_grazing | Zotu1837 | Sphingomonas | 69 | 0,315 | 0,008 |
| adaptive_grazing | Zotu922 | Azomonas | 27 | 0,222 | 0,046 |
| adaptive_grazing | Zotu1212 | Telluria | 46 | 0,21 | 0,048 |
| adaptive_grazing | Zotu218 | Pseudomonas | 100 | 0,204 | 0,036 |
| adaptive_grazing | Zotu577 | Massilia | 100 | 0,203 | 0,048 |
| all_in | Zotu860 | Pseudomonas | 100 | 0,265 | 0,019 |
| Control_adaptive_grazing | Zotu1780 | Rhodococcus | 92 | 0,302 | 0,007 |
| Control_adaptive_grazing | Zotu1181 | Hymenobacter | 100 | 0,283 | 0,005 |
| Control_adaptive_grazing | Zotu1189 | Hymenobacter | 100 | 0,275 | 0,009 |
| Control_adaptive_grazing | Zotu895 | Hymenobacter | 100 | 0,265 | 0,014 |
| Control_adaptive_grazing | Zotu1479 | Hymenobacter | 100 | 0,255 | 0,019 |
| Control_adaptive_grazing | Zotu596 | Hymenobacter | 100 | 0,254 | 0,021 |
| Control_adaptive_grazing | Zotu77 | Herbaspirillum | 56 | 0,247 | 0,045 |
| Control_adaptive_grazing | Zotu527 | Hymenobacter | 100 | 0,241 | 0,027 |
| Control_adaptive_grazing | Zotu764 | Hymenobacter | 100 | 0,241 | 0,021 |
| Control_adaptive_grazing | Zotu1054 | Hymenobacter | 100 | 0,236 | 0,05 |
| Control_adaptive_grazing | Zotu286 | Kineococcus | 100 | 0,236 | 0,021 |
| Control_adaptive_grazing | Zotu711 | Hymenobacter | 100 | 0,234 | 0,019 |
| Control_adaptive_grazing | Zotu1774 | Hymenobacter | 58 | 0,232 | 0,03 |
| Control_adaptive_grazing | Zotu72 | Methylobacterium | 100 | 0,23 | 0,004 |
| Control_adaptive_grazing | Zotu1374 | Massilia | 85 | 0,229 | 0,05 |
| Control_adaptive_grazing | Zotu200 | Lacisediminimonas | 78 | 0,226 | 0,045 |
| Control_adaptive_grazing | Zotu832 | Actinoplanes | 82 | 0,225 | 0,04 |
| Control_adaptive_grazing | Zotu1232 | Hymenobacter | 100 | 0,21 | 0,02 |
| Control_all_in | Zotu1755 | Oryzihumus | 92 | 0,312 | 0,001 |
| Control_all_in | Zotu562 | Terrabacter | 100 | 0,295 | 0,002 |
| Control_all_in | Zotu631 | Pseudomonas | 100 | 0,263 | 0,011 |
| Control_catch_crop | Zotu1250 | Pseudarthrobacter | 90 | 0,249 | 0,026 |
| Control_leymix | Zotu801 | Hymenobacter | 100 | 0,336 | 0,003 |
| Control_leymix | Zotu1833 | Pedobacter | 100 | 0,332 | 0,001 |
| Control_leymix | Zotu1043 | Herbiconiux | 87 | 0,321 | 0,001 |
| Control_leymix | Zotu758 | Variovorax | 79 | 0,314 | 0,002 |
| Control_leymix | Zotu1076 | Polaromonas | 47 | 0,308 | 0,004 |
| Control_leymix | Zotu1660 | Chryseobacterium | 100 | 0,303 | 0,001 |
| Control_leymix | Zotu1434 | Mucilaginibacter | 100 | 0,293 | 0,006 |
| Control_leymix | Zotu1392 | Chryseobacterium | 100 | 0,29 | 0,01 |
| Control_leymix | Zotu1817 | Variovorax | 41 | 0,29 | 0,013 |
| Control_leymix | Zotu1638 | Subtercola | 84 | 0,29 | 0,013 |
| Control_leymix | Zotu1413 | Hymenobacter | 100 | 0,289 | 0,013 |
| Control_leymix | Zotu1685 | Chryseobacterium | 100 | 0,286 | 0,009 |
| Control_leymix | Zotu1244 | Pedobacter | 100 | 0,283 | 0,001 |
| Control_leymix | Zotu1672 | Variovorax | 100 | 0,282 | 0,004 |
| Control_leymix | Zotu1529 | Hymenobacter | 100 | 0,268 | 0,011 |
| Control_leymix | Zotu1175 | Hymenobacter | 100 | 0,264 | 0,011 |
| Control_leymix | Zotu681 | Hymenobacter | 100 | 0,26 | 0,015 |
| Control_leymix | Zotu236 | Methylobacterium | 99 | 0,259 | 0,012 |
| Control_leymix | Zotu1170 | Simplicispira | 38 | 0,245 | 0,041 |
| Control_leymix | Zotu1514 | Salinibacterium | 65 | 0,245 | 0,013 |
| Control_leymix | Zotu1467 | Nakamurella | 100 | 0,245 | 0,038 |
| Control_leymix | Zotu1331 | Mucilaginibacter | 100 | 0,242 | 0,01 |
| Control_leymix | Zotu215 | Frondihabitans | 65 | 0,241 | 0,027 |
| Control_leymix | Zotu963 | Hymenobacter | 100 | 0,239 | 0,045 |
| Control_leymix | Zotu1423 | Hymenobacter | 100 | 0,239 | 0,026 |
| Control_leymix | Zotu614 | Hymenobacter | 100 | 0,238 | 0,003 |
| Control_leymix | Zotu962 | Pedobacter | 100 | 0,238 | 0,046 |
| Control_leymix | Zotu179 | Chryseobacterium | 100 | 0,237 | 0,027 |
| Control_leymix | Zotu881 | Nakamurella | 100 | 0,235 | 0,038 |
| Control_leymix | Zotu452 | Tardiphaga | 43 | 0,235 | 0,036 |
| Control_leymix | Zotu552 | Methylobacterium | 98 | 0,234 | 0,047 |
| Control_leymix | Zotu1642 | Hymenobacter | 74 | 0,232 | 0,04 |
| Control_leymix | Zotu885 | Pedobacter | 100 | 0,232 | 0,031 |
| Control_leymix | Zotu1426 | Hymenobacter | 100 | 0,229 | 0,025 |
| Control_leymix | Zotu139 | Rhizobium | 100 | 0,229 | 0,007 |
| Control_leymix | Zotu254 | Spirosoma | 100 | 0,228 | 0,011 |
| Control_leymix | Zotu359 | Massilia | 97 | 0,225 | 0,042 |
| Control_leymix | Zotu311 | Hymenobacter | 100 | 0,221 | 0,001 |
| Control_leymix | Zotu1176 | Pedobacter | 100 | 0,217 | 0,009 |
| Control_leymix | Zotu333 | Hymenobacter | 100 | 0,208 | 0,009 |
| leymix | Zotu121 | Sphingomonas | 100 | 0,275 | 0,006 |
| leymix | Zotu1430 | Bosea | 100 | 0,264 | 0,018 |
| leymix | Zotu1035 | Polaromonas | 59 | 0,262 | 0,019 |
| leymix | Zotu1694 | Hymenobacter | 51 | 0,261 | 0,017 |
| leymix | Zotu1601 | Sphingomonas | 100 | 0,257 | 0,005 |
| leymix | Zotu505 | Pararhizobium | 51 | 0,242 | 0,018 |
| leymix | Zotu858 | Hymenobacter | 100 | 0,214 | 0,021 |

**Table S3.** Indicator species analysis of the plant leaf bacterial communities in function of the carbon farming practices. Indicator Zotus are displayed with their taxonomic attribution according to SILVA. α = 0.05, n = 229.

| **Disease symptom** | **ZOTU** | **Fungal species** | **Accession** | **Sequence similarity** | **stat** | **p-value** |
| --- | --- | --- | --- | --- | --- | --- |
| Rust | Zotu12 | *Pleosporaceae sp.** | SH224728.06FU | 100% | 0.551 | 0.019 |
| Rust | Zotu159 | *Phoma brasiliensis** | SH202145.06FU | 95% | 0.537 | 0.034 |
| Rust | Zotu61 | *Cryptococcus sp.* | SH198056.06FU | 100% | 0.516 | 0.045 |
| Rust | Zotu43 | *Rhexocercosporidi-um panacis* | SH204719.06FU | 95% | 0.485 | 0.034 |
| Rust | Zotu93 | *Helotiales sp.* | SH191096.06FU | 99% | 0.483 | 0.037 |
| Mildew | Zotu4333 | *Cryptococcus sp.* | SH198056.06FU | 84% | 0.275 | 0.048 |
| Mildew | Zotu27 | *Ascomycota sp.* | SH227804.06FU | 100% | 0.265 | 0.038 |
| Mildew | Zotu159 | *Phoma brasiliensis** | SH202145.06FU | 95% | 0.248 | 0.032 |
| Virus | Zotu78 | *Alternaria eichhorniae** | SH224789.06FU | 100% | 0.295 | 0.048 |
| Virus | Zotu38 | *Cryptococcus victoriae* | SH198055.06FU | 100% | 0.28 | 0.044 |
| Virus | Zotu73 | *Basidiomycota sp.* | SH196706.06FU | 100% | 0.181 | 0.044 |
| Large hole | Zotu38 | *Cryptococcus victoriae* | SH198055.06FU | 100% | 0.207 | 0.002 |
| Large hole | Zotu4 | *Cryptococcus sp.* | SH198056.06FU | 100% | 0.206 | 0.021 |
| Large hole | Zotu34 | *Alternaria metachromatica** | SH224792.06FU | 100% | 0.156 | 0.02 |
| Small hole | Zotu34 | *Alternaria metachromatica** | SH224792.06FU | 100% | 0.168 | 0.017 |
| Window | Zotu147 | *Dothideomycetes sp.* | SH227809.06FU | 100% | 0.201 | 0.011 |
| Aphid | Zotu80 | *Cladosporium sp. Chiang* 1588* | SH234501.06FU | 12% | 0.191 | 0.02 |
| Aphid | Zotu41 | *Dissoconium proteae** | SH197939.06FU | 100% | 0.166 | 0.038 |
| Miner | Zotu69 | *Phaeosphaeria sp.** | SH227803.06FU | 100% | 0.329 | 0.013 |
| Miner | Zotu159 | *Phoma brasiliensis** | SH202145.06FU | 95% | 0.315 | 0.022 |
| Miner | Zotu166 | *Dioszegia hungarica* | SH196961.06FU | 100% | 0.31 | 0.029 |
| Miner | Zotu71 | *Bullera globospora* | SH197114.06FU | 98% | 0.277 | 0.042 |
| Scrape | Zotu3 | *Cryptococcus victoriae* | SH198055.06FU | 100% | 0.87 | 0.002 |
| Scrape | Zotu58 | *Tremellomycetes sp.* | SH230598.06FU | 100% | 0.605 | 0.009 |
| Scrape | Zotu69 | *Phaeosphaeria sp.** | SH227803.06FU | 100% | 0.561 | 0.011 |
| Scrape | Zotu216 | *Sporidiobolales sp.* | SH228919.06FU | 100% | 0.558 | 0.018 |
| Scrape | Zotu128 | *Cadophora luteo-olivacea* | SH204718.06FU | 97% | 0.502 | 0.04 |
| Scrape | Zotu64 | *Tremellomycetes sp.* | SH230598.06FU | 100% | 0.135 | 0.049 |
| Mite | Zotu84 | *Sphaerulina pseudovirgaureae* | SH212655.06FU | 95% | 0.352 | 0.041 |
| Gall | Zotu105 | *Stagonospora pseudovitensis** | SH199974.06FU | 57% | 0.998 | 0.011 |
| Gall | Zotu151 | *Hyaloscyphaceae sp.* | SH197899.06FU | 21% | 0.987 | 0.011 |
| Gall | Zotu19 | *Dioszegia buhagiarii* | SH196959.06FU | 100% | 0.962 | 0.016 |
| Gall | Zotu133 | *Pleosporales sp.* | SH196178.06FU | 100% | 0.902 | 0.018 |
| Gall | Zotu55 | *Ascomycota sp.* | SH206984.06FU | 100% | 0.877 | 0.042 |
| Gall | Zotu171 | *Stagonospora pseudovitensis** | SH199974.06FU | 100% | 0.864 | 0.045 |
| Gall | Zotu83 | *Oculimacula yallundae** | SH204725.06FU | 83% | 0.74 | 0.044 |
| Other | Zotu18 | *Pleosporales sp.** | SH233951.06FU | 100% | 0.571 | 0.011 |
| Other | Zotu83 | *Oculimacula yallundae** | SH204725.06FU | 83% | 0.555 | 0.022 |
| Other | Zotu93 | *Helotiales sp.* | SH191096.06FU | 99% | 0.513 | 0.042 |
| Other | Zotu121 | *Ascomycota sp.* | SH206984.06FU | 100% | 0.496 | 0.045 |
| Other | Zotu99 | *Fusarium sporotrichioides** | SH217307.06FU | 88% | 0.274 | 0.034 |

Table S4. Indicator species analysis of the plant leaf fungal communities in function of the disease symptoms present on the leaves. Indicator Zotus are displayed with their taxonomic attribution according to UNITE, potential pathogens are marked with and asterisk. α = 0.05, n = 207.

| **Disease symptom** | **ZOTU** | **Genus** | **Genus confidence (%)** | **stat** | **p-value** |
| --- | --- | --- | --- | --- | --- |
| Rust | Zotu1409 | Alpinimonas | 39 | 0.994 | 0.008 |
| Rust | Zotu142 | Frondihabitans | 57 | 0.99 | 0.008 |
| Rust | Zotu112 | Curtobacterium | 72 | 0.987 | 0.008 |
| Rust | Zotu106 | Microterricola | 67 | 0.98 | 0.008 |
| Rust | Zotu83 | Sphingomonas | 99 | 0.977 | 0.017 |
| Rust | Zotu182 | Sphingomonas | 100 | 0.976 | 0.017 |
| Rust | Zotu255 | Hymenobacter | 100 | 0.966 | 0.021 |
| Rust | Zotu1398 | Hephaestia | 17 | 0.949 | 0.008 |
| Rust | Zotu263 | Sphingomonas | 84 | 0.947 | 0.03 |
| Rust | Zotu284 | Sphingomonas | 59 | 0.947 | 0.026 |
| Rust | Zotu1166 | Hephaestia | 8 | 0.91 | 0.031 |
| Rust | Zotu119 | Methylobacterium | 100 | 0.909 | 0.035 |
| Rust | Zotu1802 | Hymenobacter | 49 | 0.894 | 0.049 |
| Rust | Zotu125 | Salinibacterium | 25 | 0.853 | 0.018 |
| Rust | Zotu91 | Methylobacterium | 99 | 0.842 | 0.03 |
| Mildew | Zotu284 | Sphingomonas | 59 | 0.334 | 0.012 |
| Mildew | Zotu182 | Sphingomonas | 100 | 0.327 | 0.019 |
| Mildew | Zotu263 | Sphingomonas | 84 | 0.321 | 0.021 |
| Mildew | Zotu83 | Sphingomonas | 99 | 0.302 | 0.021 |
| Mildew | Zotu92 | Sphingomonas | 100 | 0.278 | 0.03 |
| Mildew | Zotu99 | Aureimonas | 78 | 0.275 | 0.032 |
| Mildew | Zotu114 | Sphingomonas | 100 | 0.271 | 0.032 |
| Large hole | Zotu170 | Clavibacter | 100 | 0.155 | 0.029 |
| Thrips | Zotu315 | Duffyella (a fungus) | 33 | 0.186 | 0.004 |
| Thrips | Zotu89 | Pantoea | 75 | 0.159 | 0.015 |
| Thrips | Zotu90 | Pantoea | 99 | 0.153 | 0.002 |
| Scrape | Zotu111 | Massilia | 95 | 0.441 | 0.017 |
| Mite | Zotu1166 | Hephaestia | 8 | 0.612 | 0.034 |
| Mite | Zotu182 | Sphingomonas | 100 | 0.599 | 0.018 |
| Mite | Zotu1398 | Hephaestia | 17 | 0.577 | 0.033 |
| Mite | Zotu83 | Sphingomonas | 99 | 0.556 | 0.02 |
| Mite | Zotu284 | Sphingomonas | 59 | 0.534 | 0.027 |

Table S5. Indicator species analysis of the plant leaf bacterial communities in function of the disease symptoms present on the leaves. Indicator Zotus are displayed with their taxonomic attribution according to SILVA, α = 0.05, n = 229.

| **Treatment** | **ITS plant hosts** | **16S plant hosts** | **Carbon farming** | **Control** |
| --- | --- | --- | --- | --- |
| Cover crop | 23 | 22 | Crop caraway, pea, rye or oats. Undersown cover crop including 1-4 species of the following: *Festuca arundinacea, Elymus caninus, Lolium multiflorum, L. multiflorum ssp. westervoldicum, L. perenne, Melilotus officinalis, Phleum pratense, Trifolium repens.* Cover crop covers the soil after cropping season | Same crop - caraway, pea, rye or oats — same as in the respective carbon farming plot— grown as monoculture. No undersown cover crop. Soil bare or covered by weeds after cropping season. |
| All-in | 19 | 24 | Combination of multiple carbon-farming practices—subsoiling, no-till/direct sowing, and nutrient-fibre soil amendments—applied with organic manure and either a ley mixture or rye crop. Crop rotation. | Same crop, ley or rye but without carbon-farming practices. Crop rotation. |
| Adaptive grazing | 18 | 16 | Cattle / sheep enclosed in smaller, movable paddocks. Daily rotation cycle. Thirty-day recovery period. | Cattle / Sheep can freely roam inside a larger enclosure. Rotation cycle weeks. |
| Ley mixture | 15 | 16 | Eight or more species ley mix including typically legumes, grasses, and flowering plant species grown for five consequtive growing seasons. | Three species ley mix including typically clovers, timothy grass, and/or a fescue species grown for five consequtive growing seasons. Ley was grown to be harvested for silage twice in a growing season. |

Table S6. Description of the different carbon farming system with their amount of plant host species in both 16S and ITS datasets.
